# RSV infection does not induce EMT

**DOI:** 10.1101/2023.03.13.532506

**Authors:** Sattya N. Talukdar, Brett McGregor, Jaspreet K. Osan, Junguk Hur, Masfique Mehedi

## Abstract

Respiratory syncytial virus (RSV) infection does not cause severe disease in most of us despite suffering from multiple RSV infections in our lives. However, infants, young children, older adults, and immunocompromised patients are unfortunately vulnerable to RSV-associated severe diseases. A recent study suggested that RSV infection causes cell expansion, resulting in bronchial wall thickening *in vitro*. Whether the virus-induced changes in the lung airway resemble epithelial-mesenchymal transition (EMT) is still unknown. Here, we report that RSV does not induce EMT in three different *in vitro* lung models: the epithelial A549 cell line, primary normal human bronchial epithelial cells, and pseudostratified airway epithelium. We found that RSV increases the cell surface area and perimeter in the infected airway epithelium, which is distinct from the effects of a potent EMT inducer, TGF-β1-driven cell elongation—indicative of cell motility. A genome-wide transcriptome analysis revealed that both RSV and TGF-β1 have distinct modulation patterns of the transcriptome, which suggests that RSV-induced changes are distinct from EMT.

## Importance

We have previously shown that RSV infects ciliated cells at the apical side of the lung airway. RSV-induced cytoskeletal inflammation contributes to an uneven increase in the height of the airway epithelium, resembling noncanonical bronchial wall thickening. RSV infection changes epithelial cell morphology by modulating actin-protein 2/3 complex-driven actin polymerization. Therefore, it is prudent to investigate whether RSV-induced cell morphological changes contribute to EMT. Our data indicate that RSV does not induce EMT in at least three different epithelial *in vitro* models: an epithelial cell line, primary epithelial cells, and pseudostratified bronchial airway epithelium.

## Introduction

RSV is a nonsegmented, negative-sense RNA virus belonging to the *Pneumoviridae* family and *Orthopneumovirus* genus that infects the upper respiratory tract and often leads to severe lower respiratory tract diseases, e.g., bronchiolitis and pneumonia (1-6). Almost every child is infected by RSV within 2 years of age and often requires hospitalization. Particularly in infants (<1 year), the prevalence of RSV infection is 16 times higher than that of influenza (7, 8). RSV is the most common pediatric pathogen in newborns, infants, and children under 6 years of age. Along with infants and children, elderly people, who have chronic lung complications (e.g., asthma and chronic obstructive pulmonary disease, COPD), cardiopulmonary complications or are immunocompromised, are also known to be most vulnerable to RSV-mediated disease exacerbation (9-11). Adults have no symptoms or self-resolving flu-like symptoms from RSV infections, despite having multiple RSV infections. Indeed, Talukder et al. recently showed that the airway epithelium of healthy adults is resilient to RSV infection (12). These researchers also showed that RSV infection causes cytoskeletal inflammation that resembles noncanonical bronchial wall thickening, which may explain how epithelial cells contribute to RSV-induced bronchiolitis in infants with a lack of a mature immune system. We conclude that RSV-induced bronchial wall thickening is a common pathophysiology, but infants particularly suffer the most from severe pathophysiology due to the thinner bronchial airways (13-15). Here, we wanted to answer the important question of whether RSV-induced cytoskeletal modulation induces EMT.

EMT is a cell remodeling process converting an epithelial cell to a mesenchymal cell and is essential for embryonic development and organ formation; however, EMT can also be stimulated in response to epithelial stress or injury (e.g., viral infection), leading to severe organ degeneration (e.g., fibrosis) as well as cancer progression (16-18). Generally, EMT induces cell migration, cell invasion, cytoskeletal reorganization, and apoptotic resistance (16). EMT can be activated by numerous pathways separately or jointly, including the transforming growth factor (TGF) superfamily (such as BMP and Nodal), fibroblast growth factor (FGF), epidermal growth factor (EGF), insulin-like growth factor-2 (IGF-II), hedgehog (HH), Wnt/β-catenin, integrin, and NF-κB pathways (19-23). Multiple studies have demonstrated that respiratory viral infection can induce significant TGF-β1 levels in bronchial and alveolar epithelial cells and that TGF-β1 acts as the central regulator of the pathogenesis of pulmonary fibrosis (24-26). TGF-β1 is a pleiotropic cytokine and potent growth factor that regulates cell proliferation, organization, differentiation, and apoptosis (27, 28). TGF-β1 can induce EMT by Smad-dependent or Smad-independent pathways and plays a critical role in carcinogenesis and fibrogenesis (29, 30). Miettinen *et al.* first showed TGF-β1 as the inducer of EMT, which is now considered the “master switch” of EMT both *in vitro* and *in vivo* (31-33).

Multiple animal models, including ferrets, calves, sheep, and rodents (rats, cotton rats, mice, guinea pigs, and hamsters), have been developed for RSV research; however, to date, no animal model has been able to recapitulate RSV pathophysiology (34-36). Instead, A549 cells, a human alveolar epithelial basal cell line derived from lung adenocarcinoma tissue, have been used as a well-studied model for respiratory research. Primary cells are preferable models compared to A549 cells because of their cancerous cell origin as well as their lack of proper epithelial properties (37). Therefore, primary epithelial cells are considered a more relevant model for understanding any RSV-EMT relationship.

In this study, primary cells (bronchiolar epithelial cells) were grown in both monolayer and air-liquid interface (ALI) cultures. ALI culture provides the well-differentiated pseudostratified mucociliary epithelium, which mostly simulates *in vivo* epithelium; therefore, ALI culture is considered a more appropriate model than a monolayer culture to study respiratory epithelial biology (38-40). We investigated whether RSV induces EMT in three different *in vitro* models. To determine EMT induction in response to RSV infection, we used TGF-β1 as a positive control and observed the expression of the epithelial marker E-cadherin and the mesenchymal markers Vimentin and α-smooth muscle actin (α-SMA) in A549 cells and primary bronchial epithelial cells. Vimentin is a type 3 intermediate filament that is considered a canonical EMT marker responsible for cell migration, cell invasion, and tumorigenesis, and its higher expression is observed in several cancers, including lung cancer (41-45). Vimentin is also involved in apoptosis and degraded by caspase-3, 6, and 9, resulting in cytoskeletal disruption, which is considered the basis of epithelial morphological changes (46, 47). α-Smooth muscle actin (α-SMA) is another predominant EMT marker and is an actin isoform responsible for fibrogenesis (48, 49). In contrast, E-cadherin is a transmembrane protein responsible for the formation of epithelial cell‒cell adhesion, which becomes less functional as a sign of EMT (50). As EMT modulates epithelial morphology, we determined multiple epithelial cell shape parameters, such as cell surface area, cell perimeter, circularity, aspect ratio and caliper diameter, because of RSV infection and TGF-β1 treatment. In addition, we compared the expression of genes linked with EMT after RSV infection and TGF-β1 treatment using RNA-seq analysis.

## Results

### RSV infection does not induce EMT in a lung epithelial cell line (A549)

To determine whether RSV induces EMT in A549 cells, we detected both mesenchymal markers (e.g., vimentin) and epithelial markers (e.g., E-cadherin) by two independent techniques: Western blotting and confocal microscopy. For the control, we treated A549 cells with TGF-β1 (10 ng/ml), which is a potent EMT inducer (51). We found that TGF-β1 treatment significantly increased total vimentin expression at 48 hours post-treatment (HPT), which is in line with a previous report (Fig. 1A and 1B) (52). In contrast, RSV-WT (MOI = 0.1) induced substantially lower vimentin expression than TGF-β1 (Fig. 1A and 1B). An obvious difference in total E-cadherin expression between RSV-infected and TGF-β1-treated A549 cells was observed. There was substantially low E-cadherin expression at 48 hours post-TGF-β1 treatment (Fig. 1C and 1D), and a similar result was observed in a previous report demonstrating TGF-β1-induced EMT in A549 cells (53). In contrast, RSV infection increased E-cadherin expression in A549 cells (Fig. 1C and 1D). Kaltenborn *et al.* reported similar results (54). Although vimentin and E-cadherin are the most common EMT markers, α-SMA is also known as an EMT marker due to its upregulation during EMT *in vitro* (55). We then studied the impact of α-SMA (42 kDa) in RSV-WT-infected cells. α-SMA expression was not elevated in the infected cells but increased due to TGF-β1 treatment (Supplementary Fig. S1A and B). We also validated vimentin and E-cadherin expression in A549 cells using immunofluorescence-based detection under a microscope (Fig. 1E and 1F). We found that vimentin expression clustered in the cytoplasm, closer to the nucleus, and was more likely associated with ER (Fig. 1E top panel). Vimentin was distributed throughout the cytoplasm in the TGF-β1-treated cells at 48 HPT (Fig. 1E, middle panel). The vimentin distribution in the RSV-WT-infected cells was similar to that in the mock-infected A549 cells (Fig. 1E, bottom panel), which confirmed our findings that there was no increase in vimentin expression in infected cells (Fig. 1A and B). Likewise, we found that TGF-β1 treatment substantially reduced E-cadherin expression, contrasting with the lack of obvious changes in expression or distribution in the mock-or RSV-infected A549 cells (Fig. 1F). These results suggest that RSV infection did not induce EMT in A549 cells.

**Fig. 1.**
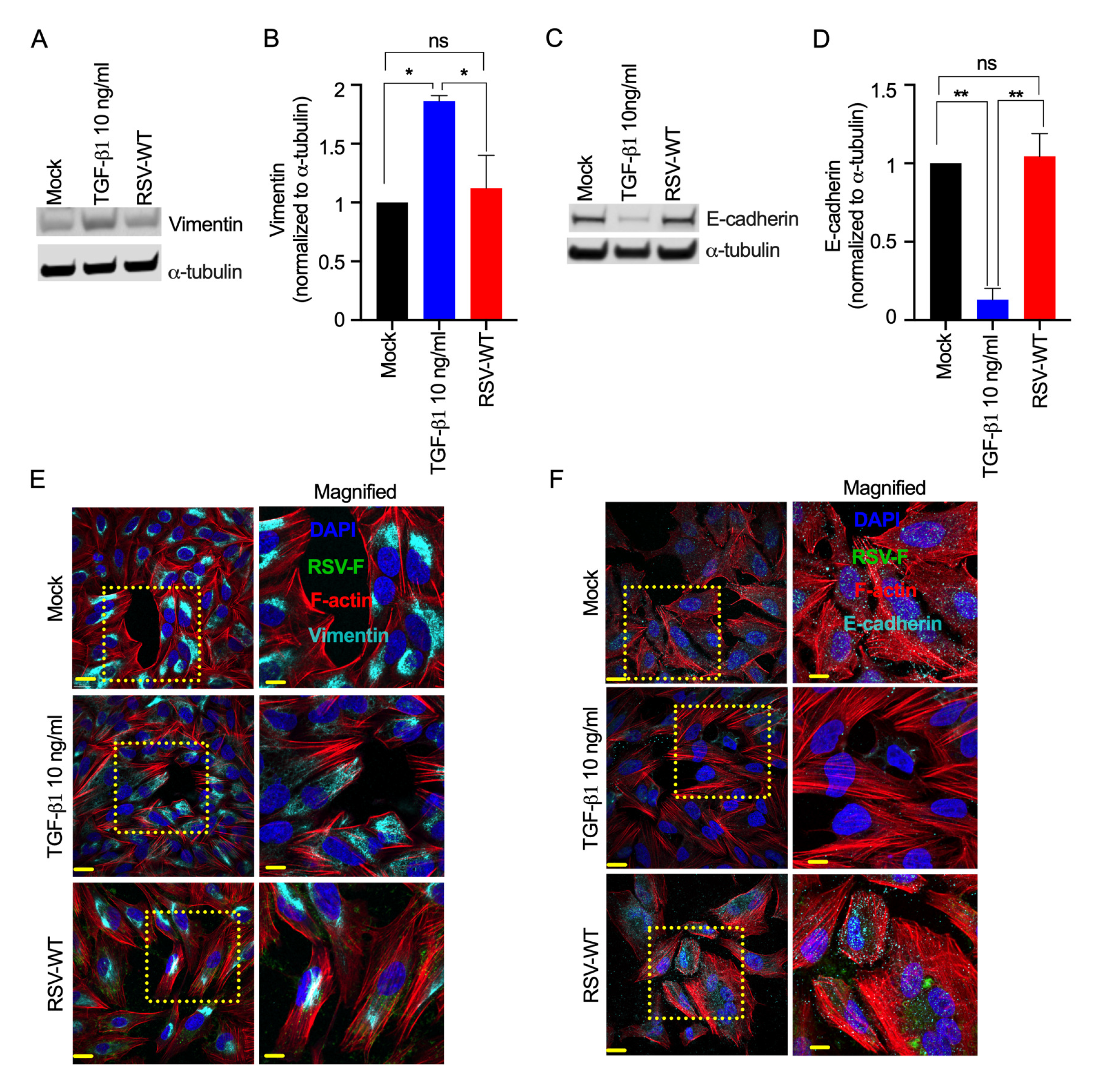
RSV does not induce EMT in A549 cells. A549 cells were mock-infected or infected with RSV-WT (MOI = 0.1) for 2 days. Separately, A549 cells were treated with TGFβ1 (10 ng/mL) as a control. **(A)** Ten micrograms of total protein was run on a reducing 4% bis-tris gel. Vimentin was detected by Western blotting using a vimentin-specific primary antibody and corresponding secondary antibody. Similarly, α-tubulin was also detected as a loading control. **(B)** Relative quantification of total vimentin (normalized to α-tubulin) in A549 cells. The data were obtained by combining the results from three independent experiments. Error bars represent the standard error of the mean (SEM). One-way ANOVA was performed to determine statistical significance. **(C).** Similarly, E-cadherin was detected. **(D)** Relative quantification of total E-cadherin (normalized to α-tubulin) in A549 cells. The data were obtained by combining the results from three independent experiments. Error bars represent SEM. One-way ANOVA was performed to determine statistical significance. **(E)** The cells were fixed, permeabilized, and immunostained for RSV F (47) and vimentin (cyan) using the respective primary antibodies and corresponding fluorescence-labeled secondary antibodies. F-actin and nuclei were visualized by rhodamine phalloidin (red) and DAPI (blue) staining, respectively. **(F)** Similarly, E-cadherin (cyan) was detected. Images were captured with a 60X objective. The scale bar is 20 µm. The scale bar for zoomed-in images is 10 µm.

### RSV infection does not induce EMT in primary normal human bronchial epithelial (NHBE) cells

To determine whether RSV infection induces EMT in primary bronchial epithelial cells, we infected NHBE cells from a healthy adult with RSV-WT (MOI = 0.1) for 2 days. For the control, we treated NHBE cells with TGF-β1 (10 ng/ml). We found that TGF-β1 treatment increased vimentin expression in NHBE cells, which suggested that TGF-β1 induced EMT in primary NHBE cells (Fig. 2A and B). Previous studies have also shown that TGF-β1 induces EMT in primary NHBE cells (49, 56). Compared to that of the TGF-β1 treatment, total vimentin expression was substantially lower in the RSV-infected primary NHBE cells (Fig. 2A and B), which suggested that RSV infection lowers or does not modulate vimentin expression. Likewise, we found that E-cadherin expression in NHBE cells was not decreased, indicating that there was no EMT in the RSV-infected NHBE cells (Fig. 2C and D). As expected, RSV infection did not increase total α-SMA expression (Fig. S2A and B). Unexpectedly, TGF-β1 treatment neither decreased total E-cadherin nor increased total α-SMA expression in the NHBE cells up to 48 hr. Although RSV-WT did not change total vimentin and E-cadherin levels in NHBE cells compared to mock infected cells, we observed a difference in their intracellular spatial distributions. We observed a higher aggregation of both vimentin and E-cadherin compared to the mock-infection and TGF-β1 treatment. While vimentin aggregated close to the nucleus, E-cadherin aggregated close to the cell-to-cell junctions (Fig. 2E and F). Whether these protein aggregations were due to RSV-induced syncytium formation in the infected cells remains to be determined. Overall, these results suggest that RSV infection did not induce EMT but rather modulated the spatial distribution of common EMT markers in the infected NHBE cells.

**Fig. 2.**
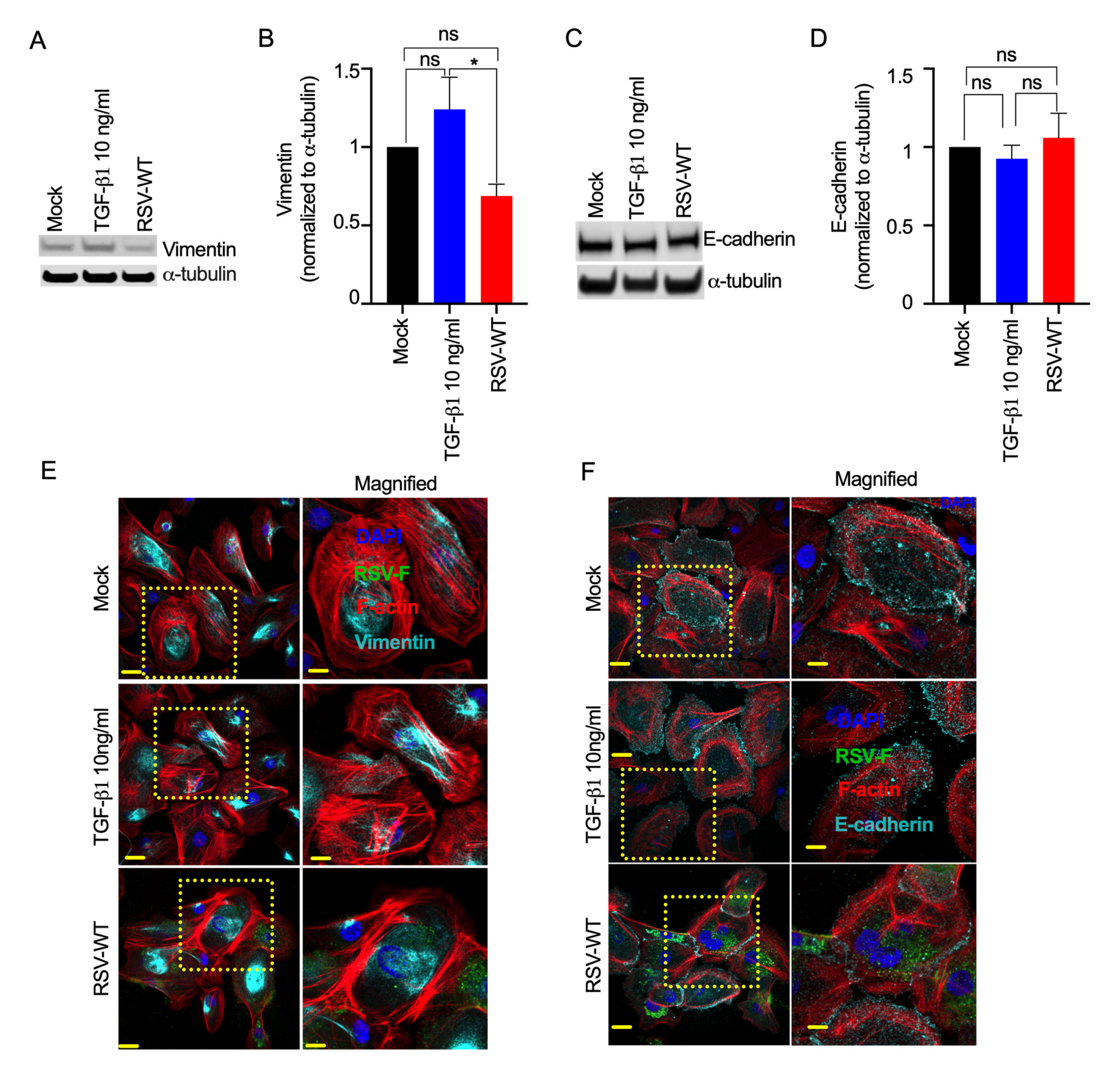
RSV infection does not induce EMT in primary epithelial cells. NHBE cells were mock-infected or infected with RSV-WT (MOI=0.1) for 2 days. Separately, NHBE cells were treated with TGFβ1 (10 ng/mL) as a control. **(A)** Ten micrograms of total protein was run on a reducing 4% bis-tris gel. Vimentin was detected by Western blotting using a vimentin-specific primary antibody and corresponding secondary antibody. Similarly, α-tubulin was also detected as a loading control. **(B)** Relative quantification of total vimentin (normalized to α-tubulin) in NHBE cells. The data were obtained by combining results from three independent experiments, and the error bars represent SEM. One-way ANOVA was performed to determine statistical significance. **(C)** Similarly, E-cadherin was detected. **(D)** Relative quantification of total E-cadherin (normalized to α-tubulin) in NHBE cells. The data were obtained by combining three independent experiments, and the error bars represent SEM. One-way ANOVA was performed to determine statistical significance. **(E)** The cells were fixed, permeabilized, and immunostained for RSV F and Vimentin (cyan) using the respective primary antibodies and corresponding fluorescence-labeled secondary antibodies. F-actin and nuclei were visualized by rhodamine phalloidin (red) and DAPI (blue) staining, respectively. **(F)** Similarly, E-cadherin (cyan) was detected. Images were captured with a 60X objective. The scale bar is 20 µm. The scale bar for magnified images is 10 µm.

### RSV infection does not induce EMT in the bronchial airway epithelium

RSV infection did not induce EMT in either a lung epithelial transformed cell line (A549) or primary NHBE cells. As both primary NHBE cells and epithelial A549 cells were grown in 2D culture, we wanted to investigate whether RSV infection induces EMT in an appropriate *in vitro* airway epithelium model that mimics the bronchial airway *in vivo*. Thus, we used a pseudostratified airway epithelium obtained by differentiating healthy adult NHBE cells in air-liquid interface (ALI) culture for 28 days following published protocols (57, 58). The airway epithelium mimics the *in vivo* lung bronchial airway epithelium and contains different epithelial cells: ciliated cells (stained with acetyl-α-tubulin), goblet cells (stained with MUC5AC), secretory cells (stained with MUC5B) (Fig. S3), and basal cells (not identified) (59-61). Importantly, the presence of tight, adherent and tricellular junctions of the airway epithelium was also confirmed by the detection of barrier-specific markers zonula occludens 1 (ZO-1), E-cadherin and tricellulin (or MARVELD2), respectively (62, 63) (Fig. S3). We also determined the optimal level of different biophysical properties (e.g., membrane permeability and ciliary function) of the airway epithelium by measuring transepithelial electrical resistance (TEER) and cilia beating frequency (CBF) as described previously (57, 58).

To determine whether RSV infection induces EMT in the bronchial airway epithelium, we infected airway epithelial cells collected from two independent healthy adult donors with RSV-expressing GFP (RSV-GFP) (MOI = 4) for 6 days. For the control, the epithelium was either mock-treated or treated with TGF-β1 (10 ng/ml) for the same duration with a daily partial change of basal medium containing TGF-β1 (10 ng/ml) where applicable (64). We found that TGF-β1 treatment induced a substantial increase in the potent EMT marker vimentin, which was detected at 6 days post-treatment (Fig. 3A). The TGF-β1-induced vimentin level was higher than that in both mock-treated and RSV-GFP-infected airway epithelia, as vimentin levels were almost undetectable in 10 µg of total protein from either mock-treated or RSV-GFP-infected airway epithelia (Fig. 3A and B). However, we detected vimentin in all samples by immunofluorescence (65) (Figs. 3C and S4). Interestingly, we found that TGF-β1 treatment modulated intracellular vimentin distribution, which appeared to be localized close to the nucleus (Figs. 3C and S4). However, the characterization of the spatial distribution of vimentin remains to be determined. As expected, we found that TGF-β1 treatment reduced E-cadherin expression in the airway epithelium (Fig. 3D and E). While TGF-β1 treatment induced EMT (high vimentin and low E-cadherin) in the bronchial airway epithelium, RSV infection neither increased vimentin nor reduced E-cadherin; rather, an increase in E-cadherin was observed (Fig. 3A, B, D, E). Although we observed donor-to-donor variation in E-cadherin expression, we could not determine the temporal regulation of E-cadherin expression in those cells. We found that RSV infection substantially increased E-cadherin levels, which contradicts the EMT phenotype (Fig. 3F). In addition to the higher level, we found abundant intracellular distribution of E-cadherin, including robust peripheral distribution in both the RSV-infected and neighboring uninfected cells (Figs. 3F and S5). This robust and abundant E-cadherin distribution appeared to be RSV specific and may suggest a novel and unique epithelial response to RSV infection (Figs. 3F and S5). Our results suggest that RSV infection does not induce classical EMT in the airway epithelium; instead, it changes infected cell morphology.

**Fig. 3.**
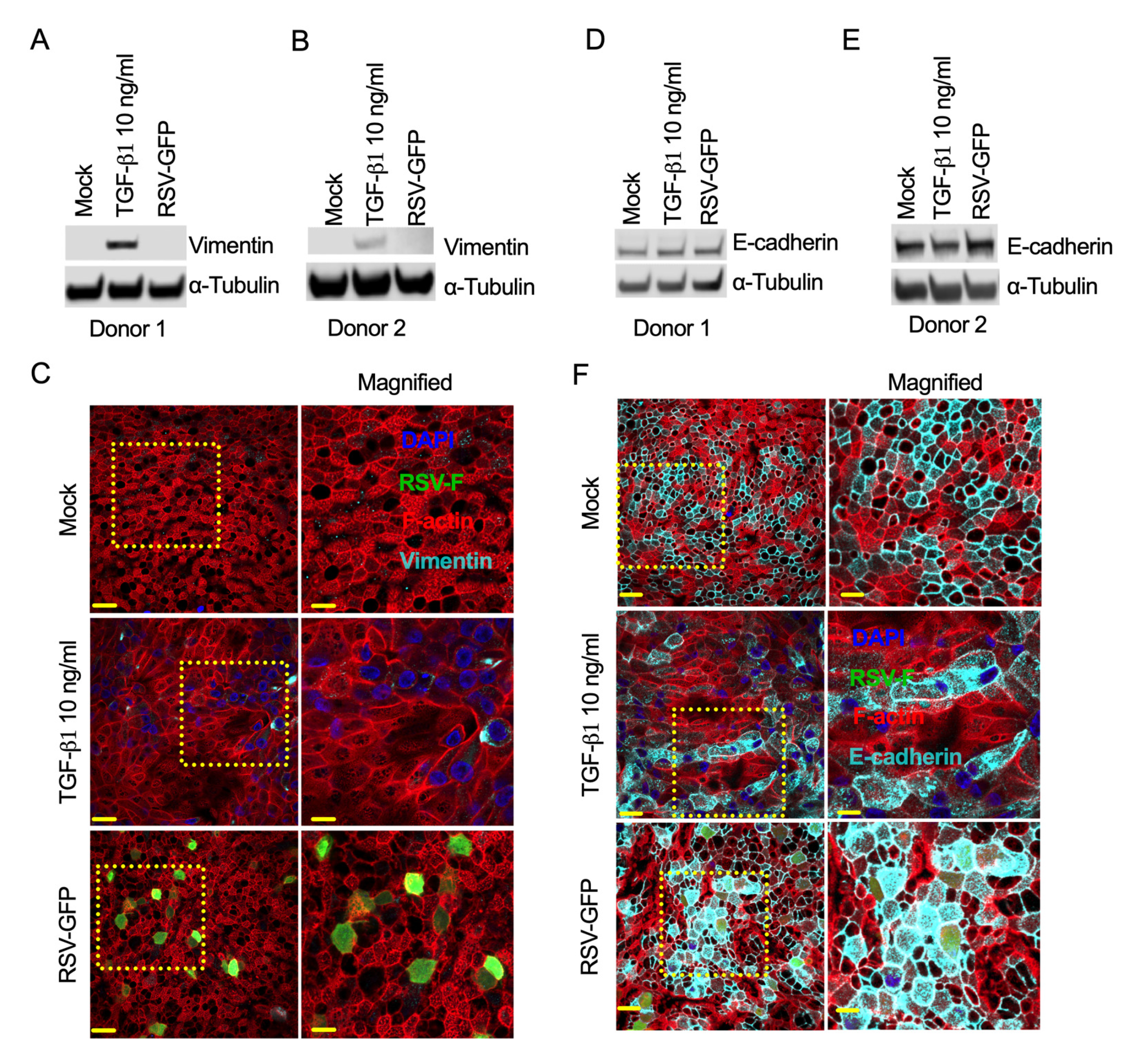
RSV infection does not induce EMT in the respiratory epithelium. The bronchial epithelium was mock-infected or infected with RSV-GFP (MOI=4) for 6 days. Separately, bronchial epithelium was treated with TGFβ1 (10 ng/mL) as a control. **(A and B)** Ten micrograms of total protein was run on a reducing 4% bis-tris gel. Vimentin was detected by Western blotting using a vimentin-specific primary antibody and corresponding secondary antibody. Similarly, α-tubulin was also detected as a loading control. The data were obtained from one independent experiment. **(C)** The cells were fixed, permeabilized, and immunostained for vimentin (cyan) using a vimentin-specific primary antibody and corresponding fluorescence-labeled secondary antibody. F-actin and nuclei were visualized by rhodamine phalloidin (red) and DAPI (blue) staining, respectively. **(D and E)** E-cadherin was also detected by Western blotting using an anti-E-cadherin primary antibody and corresponding secondary antibody. **(F)** Similarly, E-cadherin (cyan) was detected. All images were captured with a 60X objective. The scale bar is 20 µm. The scale bar for magnified images is 10 µm.

Additionally, we found similar levels of vimentin and E-cadherin in both RSV-WT-infected and RSV-GFP-infected airway epithelia. This result may confirm that RSV-GFP is a surrogate for RSV-WT (Fig. S6A and B). Moreover, we assessed the expression level of another common EMT marker, α-SMA, in the RSV-GFP-infected airway epithelium from the two independent donors. We found that RSV-GFP infection did not increase α-SMA levels compared to those in the mock-infected airway epithelium (Fig. S7A and B). Overall, our results suggested that RSV infection did not induce classical EMT. However, RSV infection induced a substantial change in the cellular morphology, which is described later.

### RSV-induced cytoskeletal expansion is a different phenotype from EMT

We found that both RSV infection and TGF-β1 treatment altered epithelial morphology, but the changes driven by RSV infection were different from those driven by TGF-β1 treatment (Fig. 4A). RSV-infected cells were substantially larger due to their expanded cytoskeleton (Figs. 4A and S8). We characterized this RSV-induced expanded cell morphology by determining several cell shape parameters, including cell area, cell perimeter, circularity, aspect ratio, and caliper or Feret diameter. We quantified these cell morphological parameters in at least 200 random cells. The cells with RSV infection were a mix of GFP-positive (indicates RSV-infected cell) and GFP-negative (indicates presumed uninfected) cells. RSV infection increased the infected-cell surface area (115.49±4.42 µm^2^), which was twofold larger than that of the mock-infected controls (52.27±1.22 µm^2^) (Fig. 4B). Interestingly, TGF-β1 treatment also increased the cell surface area more than twofold (122.50±3.23 µm^2^) over that of the mock-treated (mock-infected) cells (Fig. 4B). However, TGF-β1 treatment induced an elongated cellular morphology in contrast to the RSV-induced circular morphology (Figs. 4A and S8). As expected, RSV infection significantly increased the cell surface perimeter in the infected cells. The cell-surface enlargement was not limited to only infected cells, as the measured cell perimeter of the cells (including infected, GFP-positive and presumed uninfected, GFP-negative) (40.25±0.76 µm) was higher than the observed mock-infected cell perimeter (28.39±0.29 µm). However, TGF-β1 treatment increased the cell perimeter (48.48±0.72 µm) slightly more than RSV infection (40.25±0.76 µm) at the same time point (Fig. 4C). Circularity is also known as the “Cell shape index” or “Shape/Form factor” determined by 4π(area)/(perimeter)^2^, and its value ranges from 0 to 1, where values closer to 0 and 1 correspond to more elongated or more circular, respectively (66). Both RSV-infected cells (0.83±0.004) and uninfected cells (0.80±0.004) were more circular than TGF-β1-treated cells (0.65±0.009) (Fig. 4D). The aspect ratio (AR) was determined by the ratio of the long axis and the short axis of epithelial cells, and a higher AR indicates higher cell shape elongation. RSV-infected cells (1.41±0.01) exhibited slightly lower AR values than uninfected cells (1.51±0.02), but cell shape elongation and AR values were significantly higher after TGF-β1 treatment (2.37±0.06) (Fig. 4E). The caliper diameter, also known as the Feret diameter, was used to determine the longest distance between two points of a specific epithelial cell. The caliper diameter was increased almost twofold after TGF-β1 treatment. RSV-infected cells also displayed a higher caliper diameter (14.82±0.28 µm) than uninfected cells (10.71±0.13 µm) but a lower value than TGF-β1-treated cells (20.47±0.37 µm) (Fig. 4F). Our results suggest that RSV does not induce EMT in the infected bronchial airway epithelium, as the RSV-induced expanded circular cell morphology is distinct from the TGF-β1-induced elongated cell morphology.

**Fig. 4.**
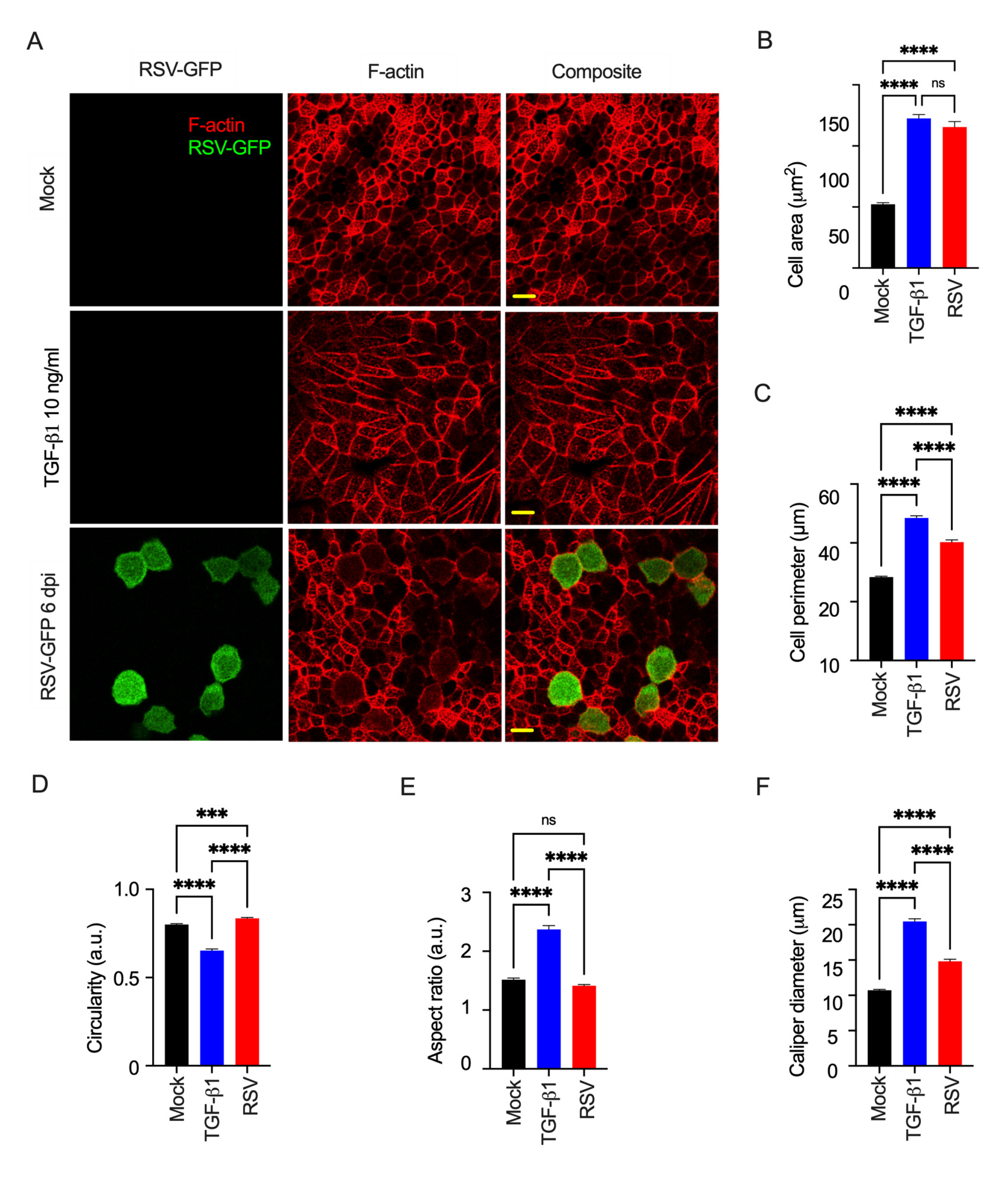
The RSV-induced epithelial morphology change is different from the TGF-β1-induced effect. **(A)** The cells from either mock-infected or mock-treated (top), TGFβ1 (10 ng/ml)-treated (middle), or RSV-GFP-infected (MOI = 4, 6 dpi) (bottom) cells were fixed, permeabilized, and stained for rhodamine phalloidin (red). Images were captured with a 60X objective. The scale bar is 10 µm. **(B-F).** Approximately 200 cells from each sample were taken for different cell shape quantifications using ImageJ software: **(B)** cell area, **(C)** cell perimeter, **(D)** circularity, **(E)** cell aspect ratio, and **(F)** caliper diameter. The data were obtained from three independent experiments. The error bars represent the standard error of the mean (SEM). One-way ANOVA was performed to determine statistical significance.

### Unique differences in the whole-genome transcriptome between RSV infection and TGF-β1 treatment

To observe the comprehensive transcriptome divergence between RSV infection and TGF-β1 treatment, we purified RNA from mock-infected, RSV-infected, or TGF-β1-treated airway epithelium and performed RNA-seq analysis. Samples were subjected to a time-stamp transcriptional analysis 6 days after RSV infection or TGF-β1 treatment (12). Comparing treatment groups to mock infection, we identified 5,863 differentially expressed genes (DEGs) as a result of TGF-β1 treatment and 4,869 DEGs influenced by RSV infection using DESeq2 with a Benjamini‒Hochberg adjusted p value <0.01 as the significance cutoff. These DEG lists were assessed for overlap, identifying 2,744 (Table S1) genes shared between RSV infection and TGF-β1 treatment as well as 2,125 (Table S2) and 3,120 genes (Table S3) unique to each group, respectively (Fig. 5A). Gene Ontology (GO) analysis was used to determine the enriched functional terms represented by genes unique to RSV infection or TGF-β1 treatment and shared DEGs within both groups. The resulting GO terms were subjected to cluster analysis to reduce redundancy in the top-represented terms; however, the full GO enrichment lists are available in the respective supplementary tables. Fig. 5B shows the top GO cluster terms for each treatment group as well as the shared DEGs. Genes unique to RSV infection represented GO terms often related to viral immune response and cellular organization (Table S4). However, genes shared between TGF-β1 treatment and RSV infection were enriched in terms related to cytoskeleton and cilium organization (Table S5). Genes uniquely induced by TGF-β1 were heavily involved in cellular migration and development (Table S6). Although the gene lists used for GO enrichment analysis were unique, some top cluster terms occurred in more than one group of genes. GO terms such as biological adhesion and intracellular signal transduction were enriched in both shared and either the TGF-β1 unique genes or RSV unique genes, respectively. Interestingly, for the shared genes between treatments that resulted in the biological adhesion enrichment results, only 30 of the 255 genes were directionally discordant between RSV and TGF-β1 (Table S7). This finding may suggest that while these functions are influenced by both RSV infection and TGF-β1 treatment, some synergistic relationships with unique genes may play a key role in the distinct responses we observed experimentally.

**Fig. 5.**
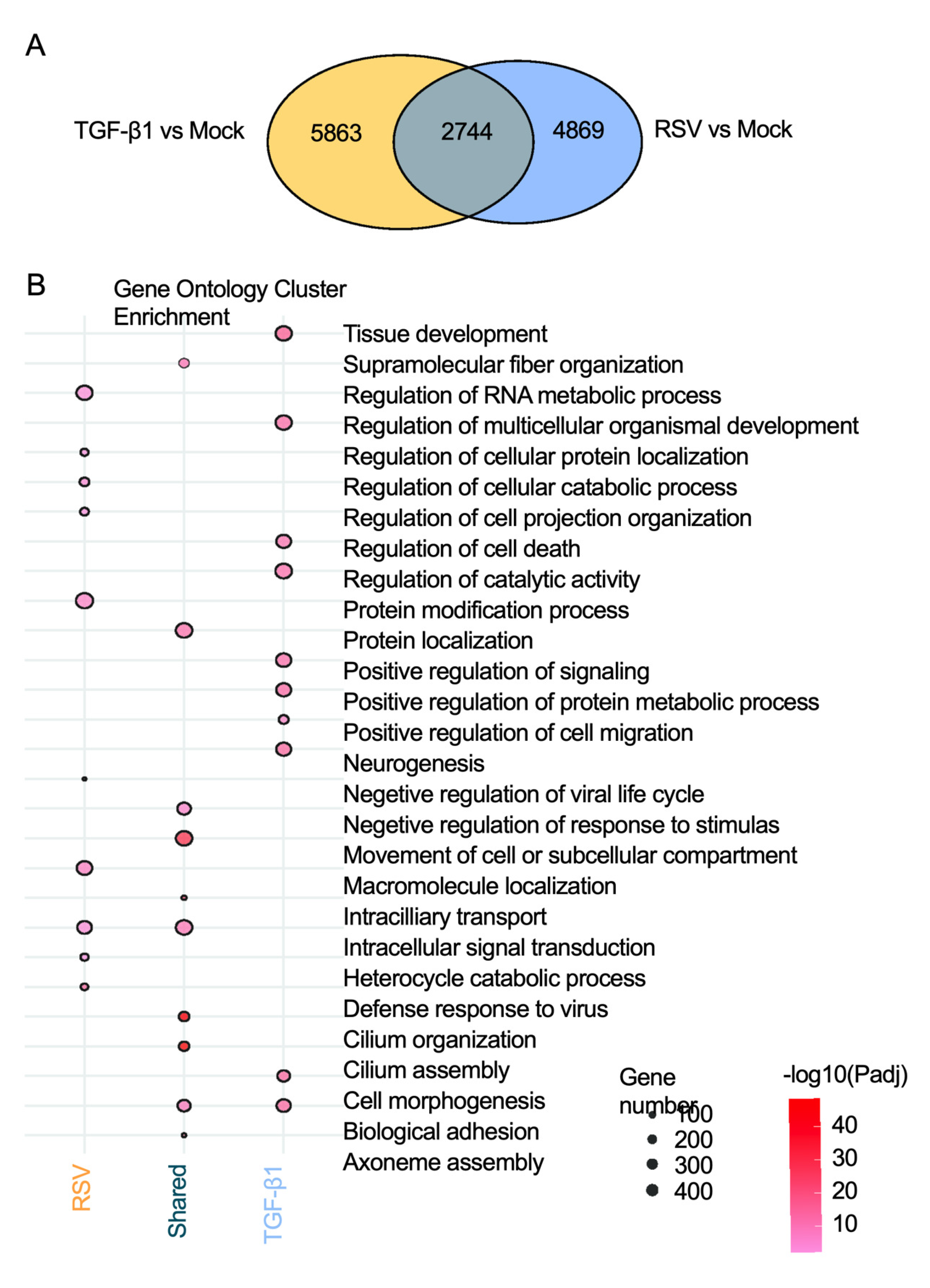
Genome-wide transcriptome differences between RSV infection and TGF-β1 treatment. **(A)** A Venn diagram represents the significant differentially expressed genes (DEGs) identified from a direct comparison between TGF-β1 vs. mock (Yellow) and RSV vs. mock (Blue) (BH adjusted p value <0.01). **(B)** A dot plot derived from Gene Ontology (GO) analysis represents the top 10 biological functions modulated by either RSV infection or TGF-β1 treatment or both. The red dots, ranging from lighter to darker shades of red, correspond to the adjusted p value for each term, with darker red indicating higher significance. The size of the dot denotes the number of genes involved in specific pathways from our supplied gene list.

### Unique differences in EMT-related gene expression between RSV infection and TGF-β1 treatment

To determine the differences in EMT-related gene expression between RSV infection and TGF-β1 treatment, we used a comprehensive list of EMT-related genes from a public database named dbEMT 2.0 (67). Out of 1,184 EMT-related human genes from dbEMT 2.0, 207 genes were commonly modulated by RSV infection and TGF-β1 treatment (Tables S8 and S9). The heatmap of the 207 EMT-related DEGs illustrates three distinctive clusters of unique expression patterns between the mock-infected, RSV-infected, and TGF-β1-treated groups (Fig. 6A). Each cluster was analyzed for GO enrichment to determine the functions represented by the varying patterns of gene expression between groups (Fig. 6B). The 34 EMT genes in Cluster 1 were mostly downregulated by both RSV infection and TGF-β1 treatment, and these genes were enriched in GO terms related to cellular development and RNA metabolism (Table S10). Cluster 2, containing 77 EMT genes, was more upregulated by TGF-β1 treatment than RSV infection, although all genes were DEGs within both comparisons to mock infection. The Cluster 2 genes represented GO terms related to morphogenesis, cell motility, and locomotion (Table S10). The 96 EMT genes included in Cluster 3 were upregulated by both RSV infection and TGF-β1 treatment; however, the pattern of gene expression was different between RSV infection and TGF-β1 treatment; specifically, RSV infection upregulated distinct genes in contrast to TGF-β1 treatment and vice versa. The Cluster 3 genes were related to GO terms involved in development, cellular proliferation/death, and cell communication (Table S10). However, within the top GO cluster terms, regulation of cell migration, other significantly enriched terms, similar to Cluster 2, such as locomotion and positive regulation of cell motility, were also significant.

**Fig. 6.**
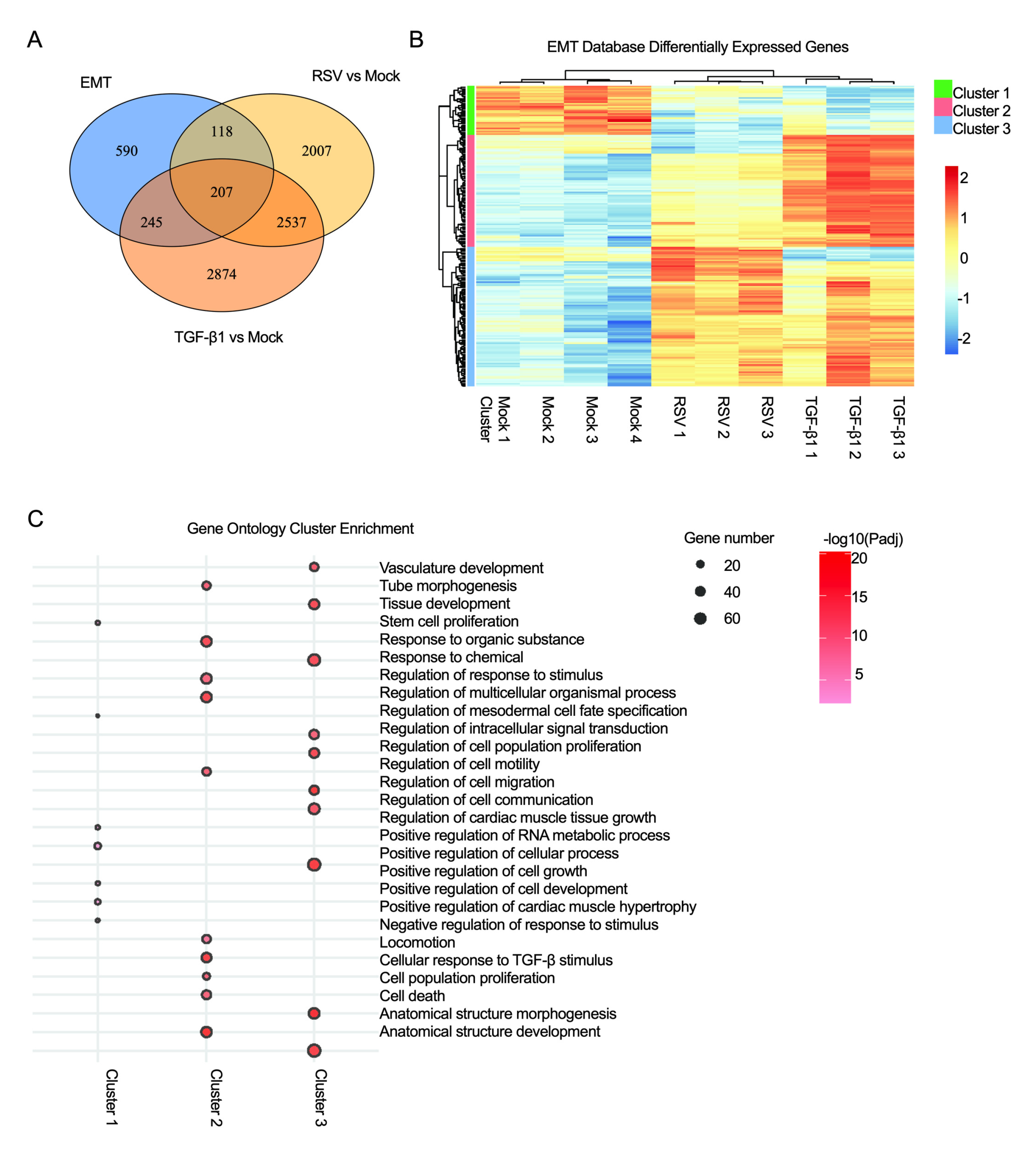
Disparities in EMT gene expression between RSV infection and TGF-β1 treatment. **(A)** A heatmap of differentially expressed EMT genes (EMT database, see methods & materials) demonstrates the comparison of 207 differentially expressed EMT-related genes between RSV infection and TGF-β1 treatment. The columns represent the samples (number indicates replicate): mock-infected or mock-treated, RSV-infected, or TGF-β1-treated. The rows represent EMT-related genes. Gene expression level is presented by pseudocolor (scale from −2 to 2); red and blue indicate the upregulation and downregulation of differentially expressed genes, respectively. Green, pink and blue bars indicate 3 different clusters of genes. **(B)** A dot plot generated by GO enrichment analysis represents the top biological functional terms modulated by EMT gene expression overlapping between RSV-infected and TGF-β1-treated samples. The GO functions presented are the top terms for each cluster of terms from each gene group. The dot size represents the number of genes, and red shading represents the adjusted p value.

## Discussion

This study aimed to investigate whether RSV induces EMT in three different *in vitro* lung models. We evaluated the expression of important EMT markers (e.g., E-cadherin, vimentin, and α-SMA) in a respiratory epithelial cell line as well as primary epithelial cells. First, we determined whether RSV induced EMT in a lung epithelial cell line (e.g., A549 cells) or primary cells (e.g., NHBE cells of healthy adults). Here, we used TGF-β1 as a positive control, as several studies have demonstrated that TGF-β1 induces EMT in A549 cells (68, 69). We used TGFβ1 (10 ng/ml) as an EMT inducer according to previous reports where a similar concentration was applied in different cell lines and primary cells grown in monolayer and ALI culture (70, 71). RSV infection did not show EMT marker alterations in our study, specifically, in human respiratory epithelial cell lines (A549 cells), but other respiratory viruses, such as rhinovirus, which is also responsible for coughing, wheezing, and shortness of breath, have been shown to induce EMT-like phenotypic and morphological changes in human bronchial epithelial cell lines (BEAS-2B cells) by lowering epithelial marker expression (72). In addition, SARS-CoV-2 induces EMT in A549 cells (ACE2 overexpressing), as observed in multiple studies (73-75). Our results also suggest no EMT induction by RSV infection in primary bronchial epithelial monolayer culture, but SARS-CoV-2-mediated EMT was observed to induce lung fibrosis as a post-COVID-19 complication in a similar model (73). Loss of E-cadherin expression and aberrant localization of vimentin expression were observed in TGF-β1-treated cells, which was also observed in a previous study, but RSV-infected cells did not show those phenotypes (76). Unlike several other respiratory viruses that cause viremia, RSV is mostly restricted to the respiratory tract, and the information regarding RSV-induced viremia and disease exacerbation is inadequate. A few previous studies have shown that RSV load in the blood can cause lung complications in a mouse model, which is a semipermissive model for RSV replication (36, 77-81). In contrast, rhinovirus has been shown to disrupt adherens and tight junctions, leading to increases in membrane permeability, which is considered a potential mechanism of rhinovirus-induced viremia and EMT (82, 83).

In addition, RSV induces epithelial morphological changes distinct from conventional epithelial structure, but this RSV-induced feature was not observed in the elongated epithelial cells induced by TGF-β1, as commonly observed in EMT (84). We found that both RSV infection and TGF-β1 treatment increased the epithelial cell area. TGF-β1-induced EMT by increasing cell area was also reported by another study (85). Our results demonstrated that TGF-β1 treatment caused a higher cell perimeter, which was also reported previously (86). We found an obvious difference between RSV infection and TGF-β1 treatment, as RSV-infected cells were substantially more circular than TGF-β1-treated cells. We also found that the expanded cytoskeleton and circularity were RSV specific. As vimentin overexpression induces EMT, Mendez et al. demonstrated that the application of exogenous vimentin by microinjection caused circularity or form factor reduction, which also supports our findings (87). The aspect ratio and caliper diameter are two other parameters used to determine cell shape elongation. TGF-β1 treatment caused a significantly higher aspect ratio and caliper diameter than those in uninfected and RSV-infected cells. ARHGAP4 is a Rho-GTPase that is important for the maintenance of the epithelial phenotype, and Kang et al. demonstrated that ARHGAP4 knockdown caused cell elongation by increasing ferret or caliper diameter and promoted EMT, which is concordant with our findings (88). Overall, the epithelial morphological analysis in our study suggests that RSV infection and TGF-β1 treatment both induced epithelial cell perimeter and cell surface area. The RSV-infected cells were more circular than the cobblestone-shaped uninfected epithelial cells, while the TGF-β1-treated cells were more elongated because of EMT induction, as reported in other studies (89-91). When there is a possibility of an RSV infection-induced increase in TGF-β1 expression (92), we have previously shown that RSV-induced TGF-β1 expression may be protective in the healthy airway epithelium (93). RSV infection in human bronchial epithelial cells induced EMT by inhibiting the type 3 interferon response; however, that study was carried out in immortalized human respiratory epithelial cells expressing hTERT and CDK4 genes without cilia beating or mucus-producing properties (94, 95). Another study demonstrated that RSV infection promotes EMT; however, that study was carried out in a monolayer culture, and the nodal gene, a member of the TGFβ family, was expressed in bronchial epithelial cells (96). There are several limitations of this study. First, there is a lack of ability to determine whether immune cells contribute to EMT in the RSV-infected airway epithelium. A lung-chip model would be helpful to determine the immune cell contribution to EMT *in vitro*, which is yet to be done. Second, there was a lack of pediatric NHBE cells to determine whether RSV induces EMT in the pediatric bronchial airway model. Third, TGF-β1-mediated EMT can be induced by either Smad-dependent or Smad-independent pathways (23). Whether RSV infection modulates Smad is yet to be determined.

The morphological analysis also supports our transcriptome analysis results in that RNA-seq demonstrated that both RSV and TGF-β1 were involved in the modulation of cell adhesion (Fig. 5B), explaining the changes in the epithelial phenotype. RSV infection did not show any specific involvement with pathways directly linked to EMT, but TGF-β1 treatment displayed its connection with EMT, as evidenced by the enriched functions related to cell migration and cellular morphogenesis (Fig. 5B). This result suggests that RSV does not induce EMT and that RSV-induced morphological changes are distinct from EMT induced by other viruses (97). Other respiratory viruses, such as influenza A and SARS-CoV-2, alter host cell morphology, but RSV-mediated epithelial changes are unique in that they do not affect membrane integrity or ciliary beating (98, 99). Furthermore, various EMT-implicated genes in the dbEST database showed a significant upregulation in genes involved in cell motility, locomotion, and anatomical structure morphogenesis upon TGF-β1 treatment; however, these genes generally remained almost unchanged after RSV infection (Fig. 6B), indicating the absence of EMT in RSV infection. While RSV-induced modulation of ARP2/3 complex-driven actin polymerization has previously been established (12, 100-103), actin polymerization modulation has only been described in EMT-induced tumor cell migration and invasion (104). The effect of RSV infection on EMT induction in COPD and pediatric models should be investigated. COPD patients have demonstrated lowered expression of E-cadherin and ZO-1 along with higher expression of vimentin (105). In addition, RSV-induced goblet cell hyperplasia/metaplasia in pediatric samples grown in ALI culture was reported (106). Goblet cell hyperplasia/metaplasia is associated with COPD; therefore, screening for EMT phenomena during RSV infection in infants and children is also important (107-109).

Overall, EMT was previously considered a simple binary state characterized by a decrease in the epithelial marker E-cadherin and an increase in the mesenchymal marker vimentin (110, 111). Later, numerous studies revealed that multiple intermediate steps are involved between epithelial and complete mesenchymal features at both morphological and transcriptional levels, which can be considered hybrid or partial EMT (112-114). Our findings, based on traditional EMT marker expression as well as morphological and transcriptome analysis, suggest that RSV infection does not cause typical EMT induction, and further investigation is warranted to determine whether RSV infection leads to partial or hybrid EMT.

## Methods

### Primary cells, cell line, and virus

Vero cells (African green monkey kidney epithelial cell line) and A549 (human alveolar epithelial basal cell line) (ATCC CCL-185) cells were obtained from Dr. Peter Collins at the National Institute of Allergy and Infectious Diseases (NIAID). Primary normal human bronchial epithelial (NHBE) cells of healthy adults were obtained from Dr. Kristina Bailey at the University of Nebraska Medical Center (UNMC), Omaha, NE, under an approved material transfer agreement (MTA) between the University of North Dakota (UND) and UNMC, Omaha, NE. The protocol for obtaining cells was reviewed by the UNMC IRB and was determined to not constitute human subjects research (#318-09-NH). RSV-WT (A2 strain), and RSV-GFP (GFP gene was inserted between the P and M genes of RSV-WT) viruses were obtained from Dr. Peter Collins at NIAID. Both viruses were grown in Vero cells and sucrose-purified using density gradient ultracentrifugation as previously published (115).

### Cell culture (A549 cells, Vero cells)

A549 cells were grown in 100 mm culture dishes (Corning, Inc.) and maintained in F-12 medium (Life Technologies) with 10% FBS, 2% penicillin/streptomycin (Thermo Fisher Scientific) and 1% amphotericin B (Thermo Fisher Scientific). Vero cells were grown in 100 mm culture dishes (Corning, Inc.) and maintained in DMEM (Sigma-Aldrich) with 5% FBS, 2% penicillin/streptomycin (Thermo Fisher Scientific) and 1% amphotericin B (Thermo Fisher Scientific).

### Primary cell culture

We used a previously described protocol (12, 58, 116). Briefly, NHBE cell monolayer passaging was also performed in a 100 mm culture dish (Corning, Inc.). The culture dish was coated with PureCol (Advanced Biometrics) before seeding cryopreserved NHBE passage zero (P0) cells, and these cryopreserved cells were thawed in a water bath. The cells were maintained in airway epithelial cell (AEC) growth medium (PromoCell) with AEC supplement (PromoCell), 2% penicillin/streptomycin (Thermo Fisher Scientific) and 1% amphotericin B (Thermo Fisher Scientific) at 37 °C in a 5% CO_2_ incubator. Cells were grown to 90% confluency with a medium change every other day. A confluent monolayer of cells was dissociated with TrypLE (Thermo Fisher Scientific), pelleted, and reseeded into a culture dish containing AEC medium with supplements for passaging. A portion of the cells was stored at - 176°C in liquid nitrogen. Cells were passaged up to four times (P4).

### Air-liquid interface (ALI) culture

We used a previously described protocol (12, 58, 116). Briefly, 6.5 mm transwells with a 0.4 µm pore polyester membrane insert (Corning, Inc.) were coated with PureCol for 20 minutes before cell seeding and then removed. NHBE cells (5×10^4^) suspended in 200 µl of AEC medium were seeded on the apical part of a transwell. Then, 500 µl of AEC medium was added to the basal part of a transwell. When cells formed a confluent layer on the transwell, the AEC medium was removed from the apical part and replaced with PneumaCult-ALI basal medium (Stemcell Technologies) with the required supplements (Stemcell Technologies), 2% penicillin/streptomycin and 1% amphotericin B in the basal part. The ALI medium was changed from the basal medium every other day. Apical surfaces were washed with 1× Dulbecco’s phosphate buffer saline (DPBS) (Thermo Fisher Scientific) once per week initially but washed more frequently when higher mucus was observed in later days. All cells were differentiated for up to four weeks (37 °C with 5% CO_2_) until the cellular and physiological properties of the epithelial layer were obtained.

### Viral infection and TGF-β1 treatment

A549 cells were grown in monolayer culture (24-well plate), and the medium was removed and washed with 1× PBS before viral infection and TGF-β1 (catalog #240-B-002, R&D Systems) treatment. RSV-WT at a multiplicity of infection of 0.1 (MOI=0.1) was mixed with OPT-MEM for one hour (37 °C with 5% CO_2_) in each well. One hour later, the medium was removed, and 500 µl of F-12 medium (Life Technologies) with 2% FBS, 2% penicillin/streptomycin (Thermo Fisher Scientific) and 1% amphotericin B (Thermo Fisher Scientific) was added. TGF-β1 (10 ng/ml) was applied by mixing with 500 µl of F-12 medium (Life Technologies) with 10% FBS, 2% penicillin/streptomycin (Thermo Fisher Scientific), and 1% amphotericin B (Thermo Fisher Scientific). The mock-treated, RSV-infected, and TGF-β1-treated cells were incubated for two days at 37 °C. NHBE cells were also grown in monolayer culture (24-well plate), and the medium was removed and washed with 1× PBS before RSV-WT infection and TGF-β1 treatment. RSV-WT (MOI=0.1) with 150 µl of AEC growth medium (PromoCell) containing supplement (PromoCell), 2% penicillin/streptomycin (Thermo Fisher Scientific) and 1% amphotericin B (Thermo Fisher Scientific) was added to each well for one hour at 37 °C in a 5% CO_2_ incubator. Then, the medium was removed, and 500 µl of AEC growth medium was replenished. TGF-β1 (10 ng/ml) was mixed with 500 µl of AEC growth medium to be applied in each well. The mock-treated, RSV-infected, and TGF-β1-treated cells in NHBE monolayer culture were incubated for two days at 37 °C. In addition, after four weeks of ALI culture, differentiated pseudostratified airway epithelium was obtained, washed with 200 µl 1× DPBS to remove mucus, and infected with RSV-WT (MOI=4) or RSV-GFP (MOI=4) for two hours (37 °C with 5% CO_2_). The virus inoculum was removed and washed 2× with 200 µl 1× DPBS. Fresh ALI medium with supplements (500 µl) was added to the basal part of the transwell, and the apical part was kept empty. A similar concentration of TGF-β1 (10 ng/ml) was mixed with ALI medium containing supplements in the basal part of the transwell. The mock-treated, RSV-infected, and TGF-β1-treated transwells were incubated for six days at 37 °C.

### Confocal microscopy

For preparation of confocal slides from A549 and NHBE cells grown in monolayer culture, both cell lines were seeded on cover glasses (24-well plate). After RSV-WT (MOI=0.1) infection and TGF-β1 treatment (10 ng/ml), two days later, cells were washed with 1× PBS, fixed with 4% paraformaldehyde (PFA) (Polysciences, Inc.) in 1× PBS for 10 minutes at room temperature, permeabilized with 0.5% Triton-X100 (Sigma‒Aldrich) for 10 minutes followed by blocking with 3% BSA solution in 1× PBS for 2 hours. After 2 washes with 1× PBS, the cells were then incubated with the following primary antibodies in 0.1% BSA solution (1X PBS) overnight in the dark at 4 °C: mouse monoclonal (1:1000) Respiratory Syncytial Virus (F-protein) (Abcam), rabbit monoclonal (1:1000) E-cadherin (Cell Signaling Technologies), and rabbit monoclonal (1:1000) Vimentin (Cell Signaling Technologies). For preparation of confocal slides from respiratory epithelium, the transwell insert was washed with 1× PBS, and both apical and basal parts were fixed with 4% PFA in 1× PBS for 30 minutes at room temperature, followed by 2× washes with 1×PBS and then blocking with 10% goat serum in immunofluorescence (IF) washing buffer (130 mM NaCl_2_, 7 mM Na_2_HPO_4_, 3.5 mM NaH_2_PO_4_, 7.7 mM NaN_3_, 0.1% BSA, 0.2% Triton-X 100 and 0.05% Tween-20) for 1 hour. After 2× washes with 1× PBS, the transwell inserts were then incubated with similar primary antibodies: E-cadherin rabbit monoclonal (1:200) (Cell Signaling Technologies) and Vimentin rabbit monoclonal (1:200) (Cell Signaling Technologies) in IF washing buffer overnight in the dark at 4 °C. For monolayer sample preparation, the day after washing with buffer, cells were incubated with secondary antibodies: anti-rabbit Alexa Fluor 647 (1:200) (Thermo Fisher Scientific) and anti-mouse Alexa Fluor 488 (1:200) in IF buffer for 3 hours in the dark at 4 °C. Then, the samples were washed 2× with IF buffer and incubated with rhodamine phalloidin (PHDR1) (1:500) (Cytoskeleton, Inc.) for 30 minutes in the dark at 4 °C. After 3× washes with IF buffer, the nuclei were stained with NucBlue Fixed Cell Stain Ready Probes (Thermo Fisher Scientific) for 30 minutes in the dark at 4 °C. Coverslips (for A549 and NHBE cells) were washed with IF buffer before mounting on microscope slides using ProLong Gold anti-fade mounting media (Life Technologies). The transwell membrane was washed with IF buffer, followed by cutting the whole membrane to set on the microscope slides (TechMed services), and similar media were used for mounting the coverslip on the slide. Images were captured using a confocal laser scanning microscope (Olympus FV3000) enabled with a 60× objective. The 405 nm laser was used to find the DAPI signal for nucleus detection, the 488 nm laser was used to activate Alexa Fluor 488 for GFP (or RSV F) detection, the 561 nm laser was used to activate rhodamine phalloidin for F-actin detection, and the 640 nm laser was used to activate Alexa Fluor 647 for E-cadherin and Vimentin detection. Imaris software version 9.5.1 (Oxford Instruments Group) was used for the conversion of Z-stack images (.oir format) to.tiff format and other image postprocessing.

### Cell shape analysis

ImageJ software was used to determine cell area, cell perimeter, circularity, aspect ratio, and caliper diameter (12). The images (.tiff format) from mock-treated, RSV-infected, and TGF-β1-treated samples were processed and converted into an 8-bit grayscale image using ImageJ. Two hundred cells were taken from each sample for quantification. Every epithelial cell was selected manually by the polygon option based on F-actin staining. RSV-infected cells were confirmed by GFP signal along with F-actin staining.

### Western blotting

Protein samples from airway epithelium were collected after 6 days of RSV infection and TGF-β1 treatment. The apical part of a transwell was washed 2× with PBS, and then, all the cells were scraped out and transferred into a 1.5 ml tube. After removal of the supernatant, cells were transferred into a QIAshredder tube and mixed with 100 µl of gel loading buffer containing 2.5 ml of 4× LDS loading buffer (Thermo Fisher Scientific), 1 proteinase inhibitor tablet, and 7.5 ml of 1× PBS. The cell mix was centrifuged at 15,000 rpm for 3 minutes. The elusion from the QIAshredder was stored at −80 °C. Protein samples from NHBE and A549 monolayers were collected after 2 days of RSV infection and TGF-β1 treatment. Cells were scraped out from 24-well plates by mixing with gel loading buffer and transferred into a QIAshredder tube followed by centrifugation at 15,000 rpm for 3 minutes. The eluate from the QIAshredder tube was stored at −80 °C. Protein concentration was measured using a BCA protein assay kit (Thermo Fisher Scientific). Protein samples were denatured at 90 °C for 10 min with 10× reducing agent (Thermo Fisher Scientific) before gel electrophoresis. Total protein ranging from 10 to 20 µg was separated on 4-12% Bis-tris SDS polyacrylamide gels, followed by dry blot transfer onto PVDF according to the manufacturer’s instructions (Life Technologies). For the loading control, mouse α-tubulin monoclonal Ab (Thermo Fisher Scientific) was used. The PVDF membranes were incubated in LI-COR blocking buffer (1:1 in 1× PBS) (LI-COR Biosciences) for 1 hour, followed by overnight incubation with primary Ab including rabbit monoclonal anti-E-cadherin (1:1000) (3195S, Cell Signaling Technology), rabbit monoclonal anti-Vimentin (1:1000) (5741, Cell Signaling Technology), and rabbit monoclonal anti-α-SMA (1:1000) (19245, Cell Signaling Technology) in blocking buffer (LI-COR Biosciences) with 1× PBS (1:1). The membranes were washed 3× for 5 minutes with 1× PBS followed by incubation with secondary antibodies including goat anti-mouse IRDye680 Ab (1:15000) (LI-COR Biosciences) and goat anti-rabbit IRDye800 Ab (1:15000) (LI-COR Biosciences) for 1 hour. After 3 washes for 5 minutes with 1× PBS, fluorescence was analyzed using the Odyssey imaging system (LI-COR Biosciences). Image Studio 5.2 software (LI-COR Biosciences) was used for densitometric analysis.

### RNA extraction

The airway epithelium cultured on 6.5 mm transwell membranes was washed and treated with RLT buffer (Qiagen) with 1% β-mercaptoethanol (Sigma‒Aldrich). Cells were scraped using a cell scrapper, collected into a QIAshredder tube and centrifuged at 15,000 rpm for 3 minutes. The eluate was used to extract total RNA using a Total DNA/RNA Extraction Kit (Qiagen) including a DNase I treatment to remove DNA in the sample according to the manufacturer’s instructions.

### RNA-seq for transcriptome analysis

RNA was extracted (as described above) from ALI cultures at 6 DPI after mock infection or infection with RSV-WT (MOI=4) or treatment with TGF-β1. [**RNA quality here in terms of RIN values**]. The sequencing library was prepared using the SMART-Seq v4 Ultra Low Input RNA Kit with nonstranded and polyA+ selection (Clontech, Takara Bio USA, Mountain View, CA, USA). Approximately 120 million 125-bp paired-end reads per sample were obtained (HiSeq 2500, Illumina, San Diego, CA, USA). RNA-seq was performed by the Genomic core, UND, in a blinded manner.

Quality assessment of the raw sequencing data was performed using FastQC v0.11.8 (117). The adapters were trimmed using Trimmomatic v0.39 (118). Cleaned reads were aligned to the human reference genome (hg19) with STAR v2.7.1a (119). Gene expression was quantified using CuffNorm v2.2.1 (120). Read counts were summarized using featureCounts v1.4.6 (121). Differentially expressed genes (DEGs) were identified in RSV-infected vs. mock-infected and TGF-β1-treated vs. mock-infected comparisons using DESeq2 v1.24.0 with a significance cutoff of <0.01 Benjamini‒Hochberg (BH)-adjusted p value (122). Enrichment analysis for DEGs was performed using our in-house R package richR (https://github.com/hurlab/richR) to identify significantly overrepresented biological functions and pathways in terms of Gene Ontology (GO) annotation. GO terms with a BH-adjusted p value of <0.05 were deemed to be significant (123). The full list of enriched GO terms was subjected to a clustering analysis to identify conceptually overlapping terms to reduce term redundancy. The most significant term from each cluster was used as the representative term.

### Statistical analysis

Parameters such as the number of independent experiments, standard error of the mean (SEM), and statistical significance are reported in the figures and figure legends. GraphPad Prism 8 software was used for statistical analysis, and where appropriate, the statistical analysis methods are noted in the figure legends. A p value less than 0.05 was considered to indicate significance. *, p<0.05; **, p<0.01, ***, p<0.001; ****, p<0.0001; ns, not significant.

### Data availability

The RNA-seq data are available through GEO under accession number GSE189537.

## Acknowledgments

This work was funded by NIH P20GM113123. We thank the Microscopy Core (UND, Grand Forks) funded by NIH P20GM103442 of the INBRE program for providing access to an Olympus FV300 confocal microscope and Dr Swojani Shrestha for technical support. We also thank the Imaging Core (UND, Grand Forks) funded by NIH P20GM113123, NIH U54GM128729, and UNDSMHS for providing access to the Imaris image analysis software. We acknowledge the Genomics core (UND, Grand Forks) for RNA-seq funded by U54GM128729 and 2P20GM104360-06A1.

---

**Fig. S1.**
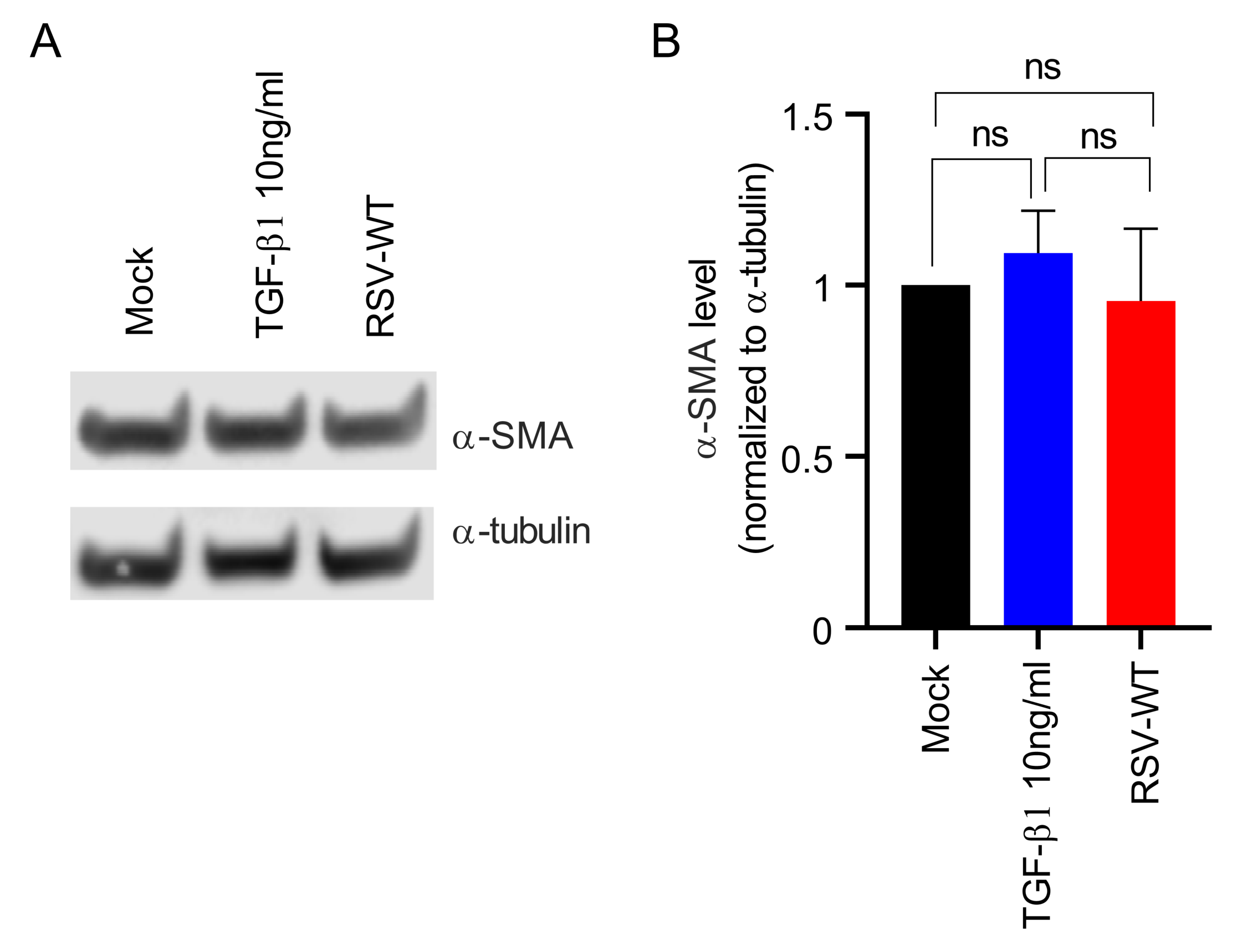
RSV infection does not increase α-SMA expression in A549 cells. A549 cells were mock-infected or infected with RSV-WT (MOI = 0.1) for 2 days. Separately, A549 cells were treated with TGFβ1 (10 ng/mL) as a control. **(A)** The cells were collected and lysed. Ten micrograms of total protein was run on a reducing 4% bis-tris gel. α-SMA was detected by Western blotting using an α-SMA-specific primary antibody and the corresponding secondary antibody. Similarly, α-tubulin was also detected as a loading control. **(B)** Relative quantification of total α-SMA (normalized to α-tubulin) in A549 cells. The data were obtained by combining results from three independent experiments, and the error bars represent SEM. One-way ANOVA was performed to determine statistical significance.

**Fig. S2.**
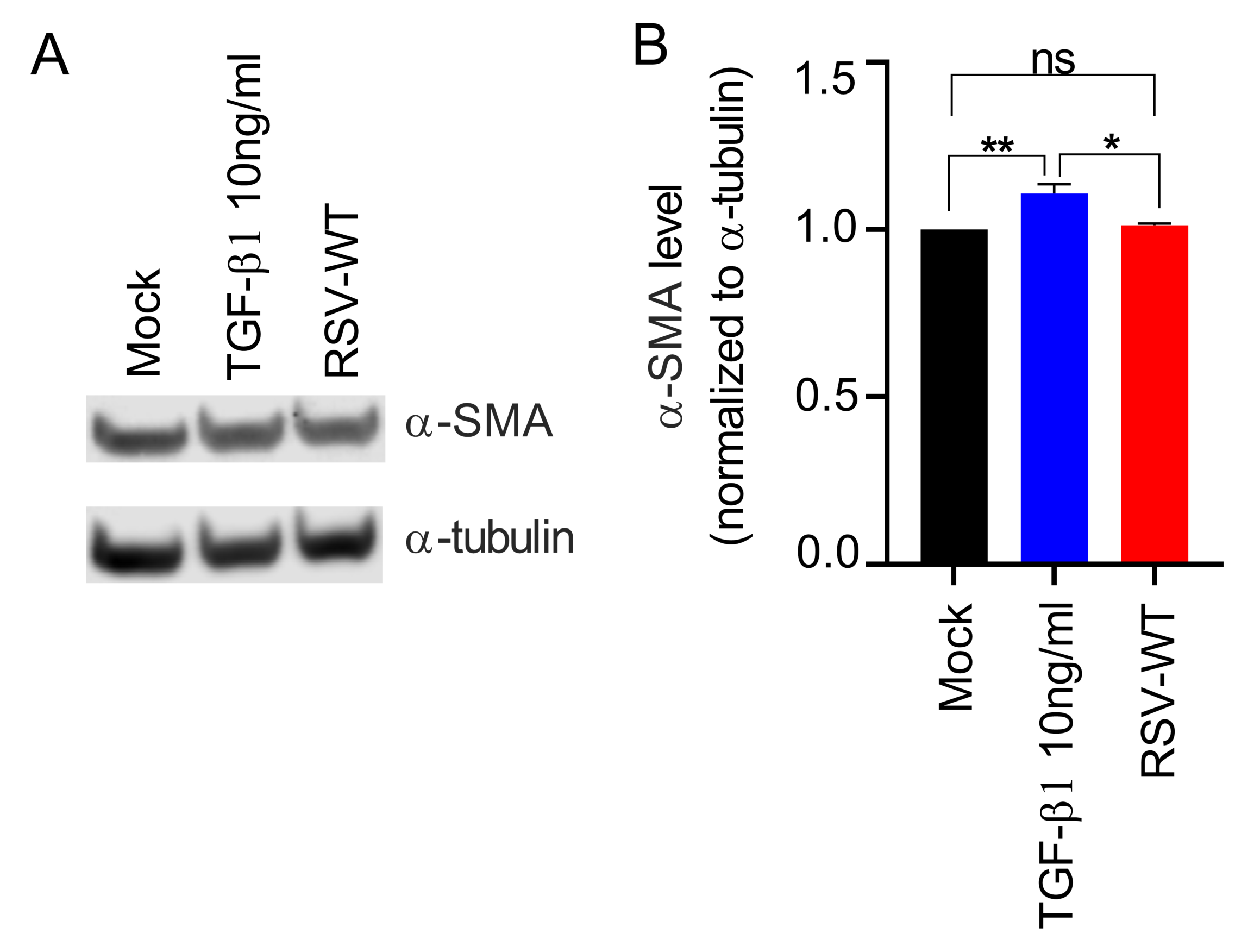
RSV infection does not increase α-SMA expression in primary epithelial cells. NHBE cells were mock-infected or infected with RSV-WT (MOI = 0.1) for 2 days. Separately, NHBE cells were treated with TGFβ1 (10 ng/mL) as a control. (A) The cells were collected and lysed. Ten micrograms of total protein was run on a reducing 4% bis-tris gel. α-SMA was detected by Western blotting using an α-SMA-specific primary antibody and the corresponding secondary antibody. Similarly, α-tubulin was also detected as a loading control. (B) Relative quantification of total α-SMA (normalized to α-tubulin) in NHBE cells. Data were obtained by combining results from three independent experiments, and the error bars represent SEM. One-way ANOVA was performed to determine statistical significance.

**Fig. S3.**
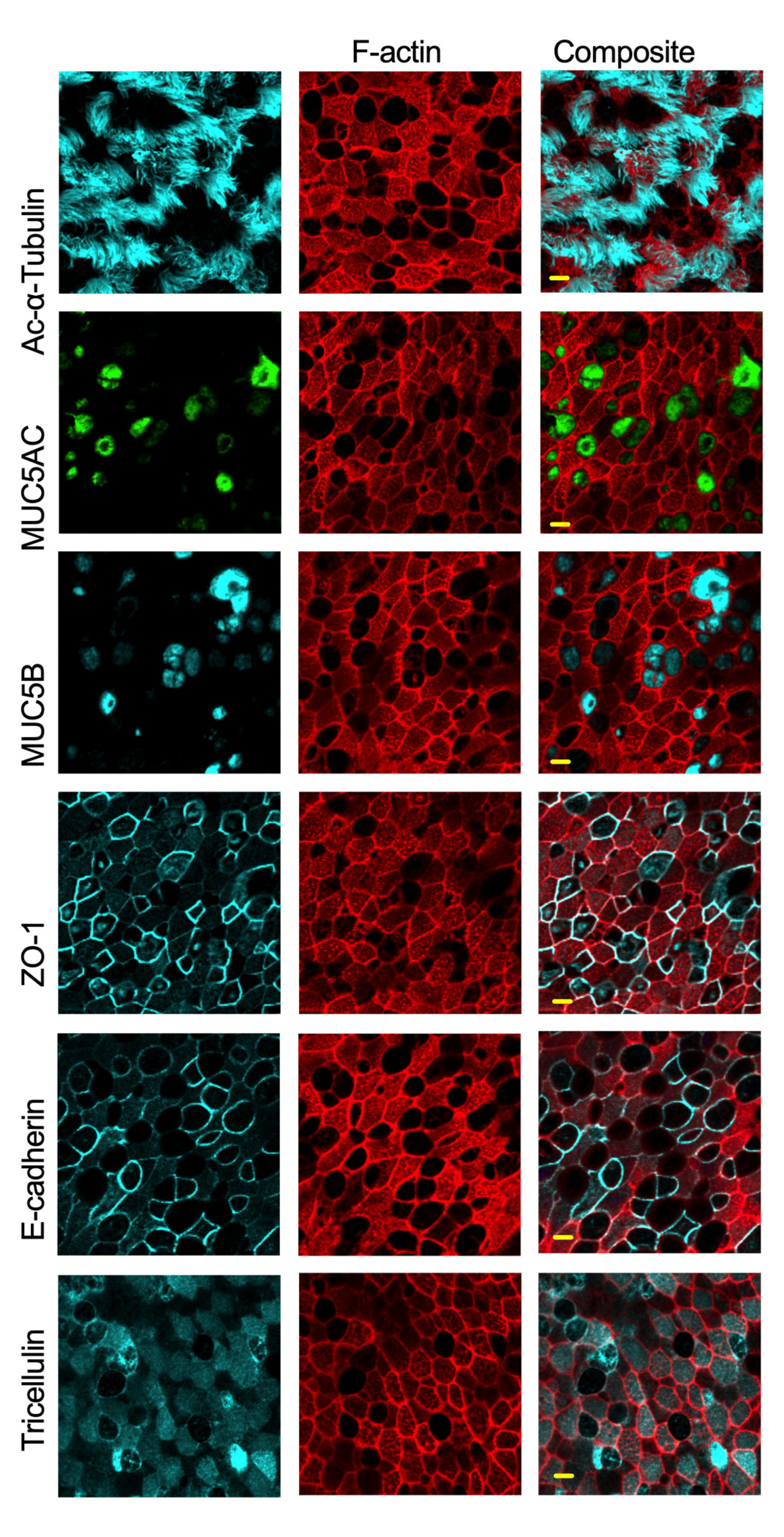
Morphology and junctional characteristics of the differentiated airway epithelium. We confirmed multicellular epithelium by detecting two important cell types. (ciliated and goblet cells) that were identified by cell-specific surface markers: for ciliated cells, we used acetyl-α-tubulin (cyan). For goblet cells, we used both MUC5AC (cyan) and MUC5B (cyan). We confirmed tissue-like airway epithelium by detecting adherens, tight, and tricellular junctions by E-cadherin (cyan), ZO-1 (cyan), and MALVELD2 (cyan) staining, respectively. The cell cytoskeleton was visualized by rhodamine phalloidin (red) staining. Images were captured with a 60× objective. The scale bar is 5 µm.

**Fig. S4.**
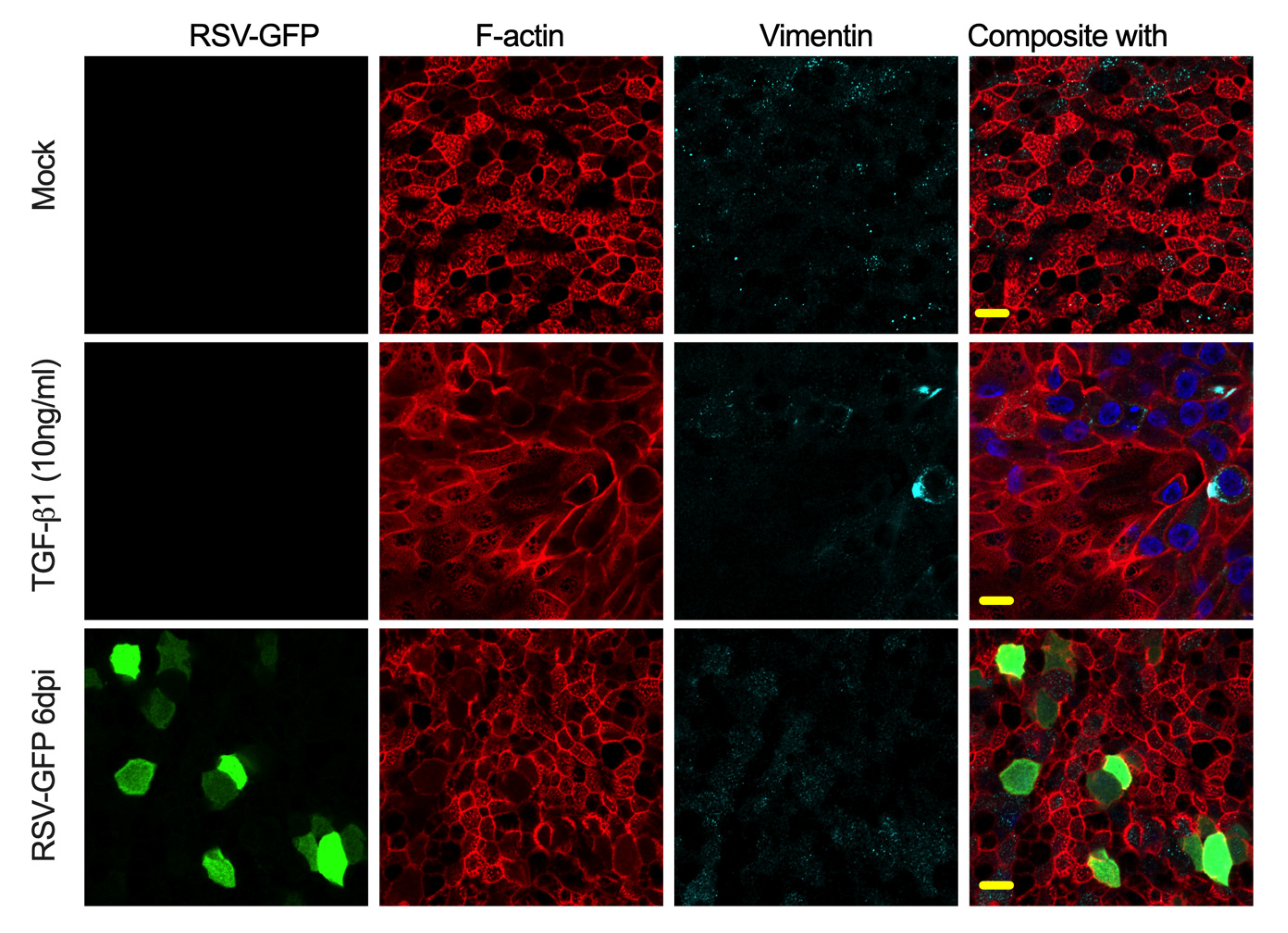
Vimentin expression determination in mock-infected or mock-treated, TGFβ1-treated or RSV-GFP-infected bronchial epithelium. The cells were fixed, permeabilized, and immunostained for vimentin (cyan) by incubating with rabbit monoclonal antibody followed by the secondary antibody anti-rabbit Alexa Fluor 647. The infected cells were identified based on the GFP signal. F-actin and nuclei were visualized by rhodamine phalloidin (red) and DAPI (blue) staining, respectively. Images were captured with a 60× objective. The scale bar is 10 µm.

**Fig. S5.**
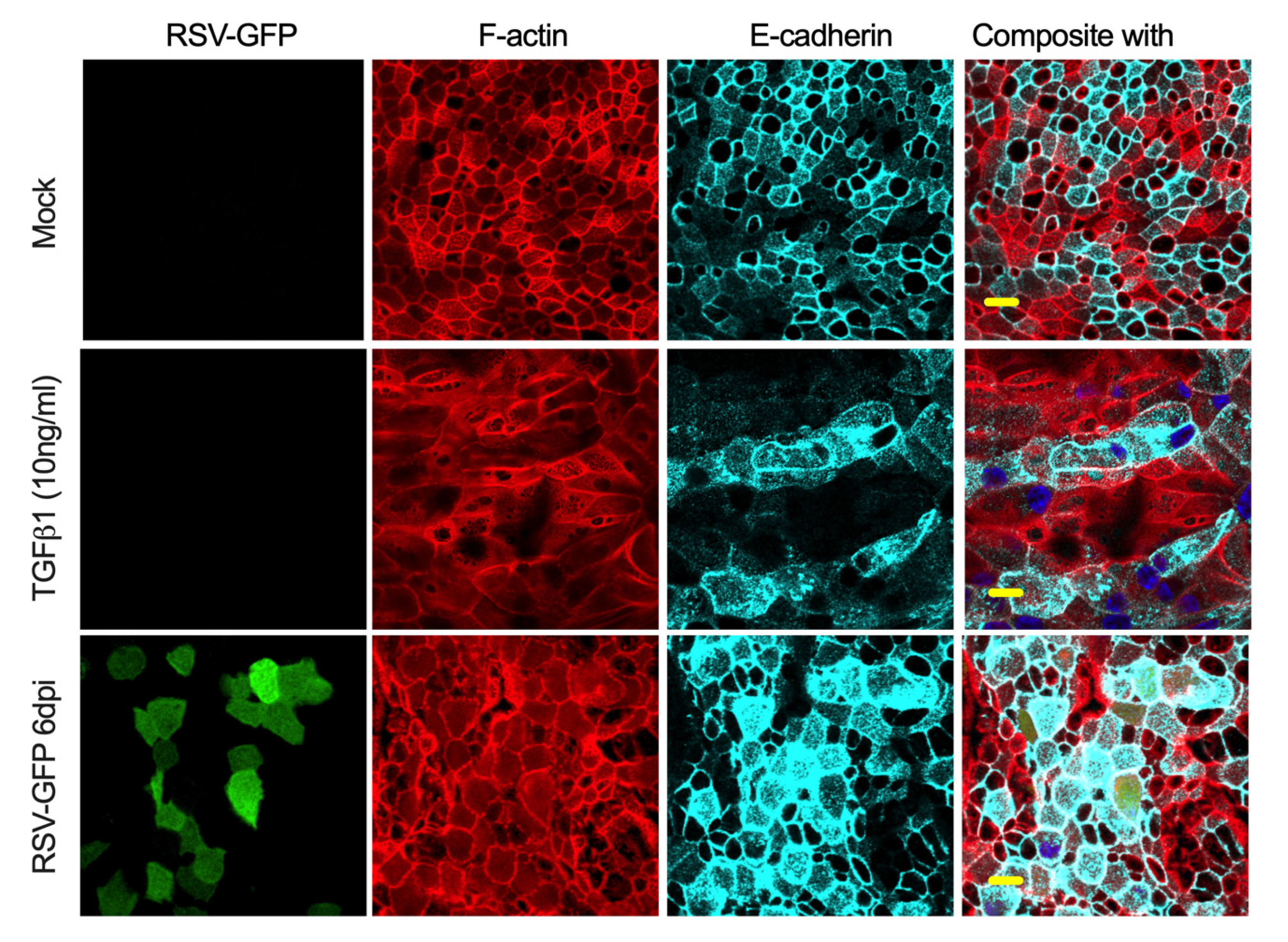
E-cadherin expression determination in mock-infected or mock-treated, TGFβ1-treated or RSV-GFP-infected bronchial epithelium. The cells were fixed, permeabilized, and immunostained for E-cadherin (cyan) by incubating with rabbit monoclonal antibody followed by secondary antibody anti-rabbit Alexa Fluor 647. The infected cells were identified based on the GFP signal. F-actin and nuclei were visualized by rhodamine phalloidin (red) and DAPI (blue) staining, respectively. Images were captured with a 60X objective. The scale bar is 10 µm.

**Fig. S6.**
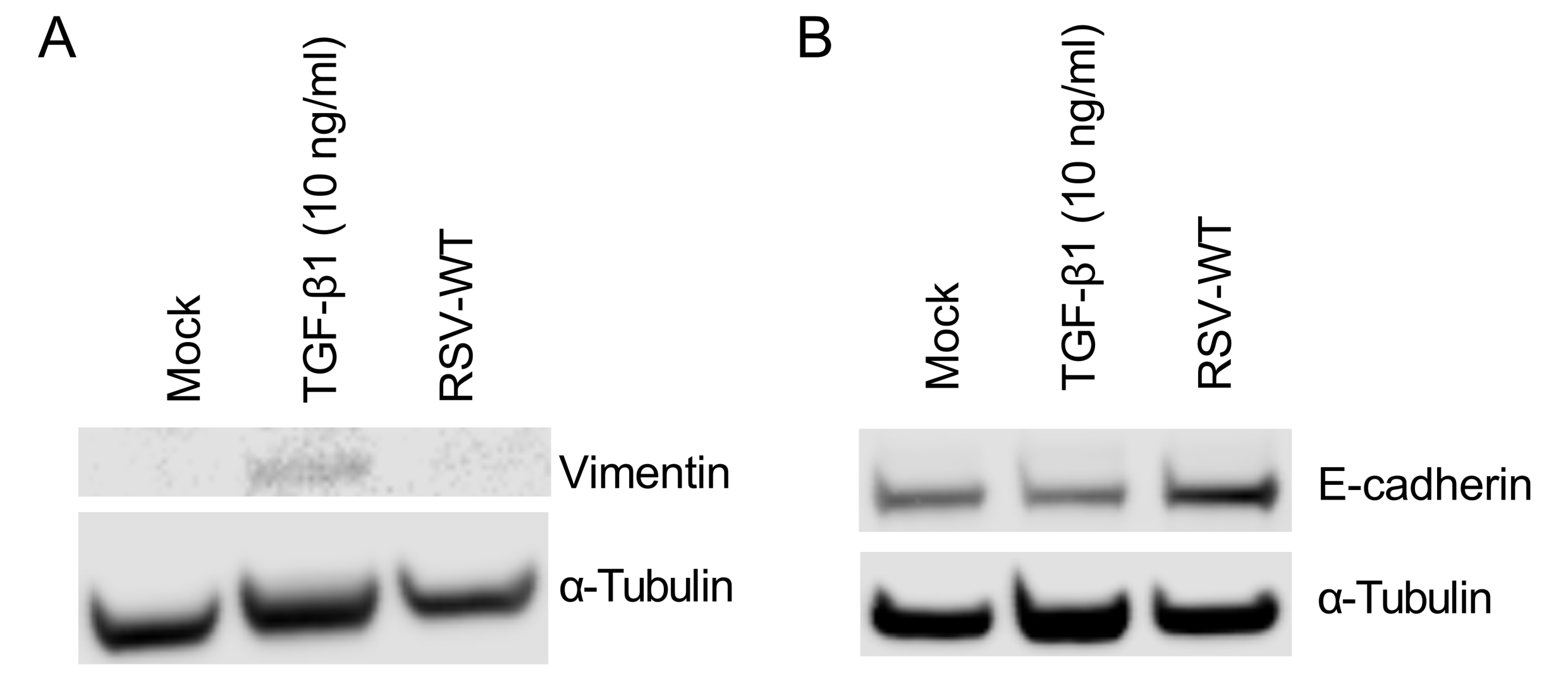
RSV-WT infection neither increased vimentin nor decreased E-cadherin expression in the respiratory epithelium. The differentiated bronchial epithelium was mock-infected or infected with RSV-WT (MOI = 4) for 6 days. Separately, the bronchial epithelium was treated with TGFβ1 (10 ng/mL) as a control. **(A)** The cells were collected and lysed. Ten micrograms of total protein was run on a reducing 4% bis-tris gel. Vimentin was detected by Western blotting using a vimentin-specific primary antibody and corresponding secondary antibody. Similarly, α-tubulin was also detected as a loading control. **(B)** E-cadherin was similarly detected. The data represent one independent experiment.

**Fig. S7.**
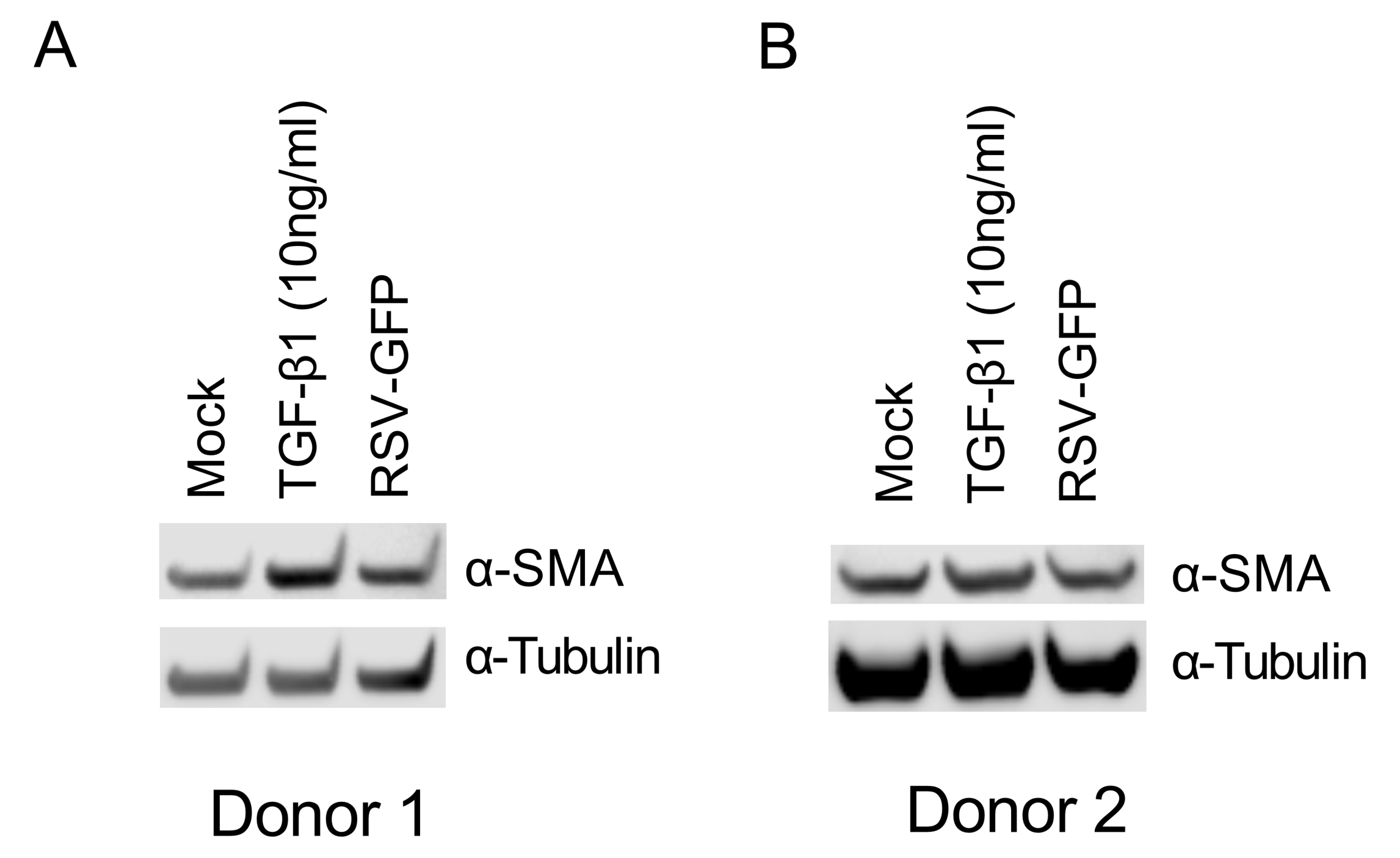
RSV infection does not increase α-SMA expression in the respiratory epithelium. The differentiated bronchial airway epithelium (Donor 1, NHBE C16 and Donor 2, NHBE E16) was mock-infected or infected with RSV-GFP (MOI = 4) for 6 days. Separately, the epithelium was treated with TGFβ1 (10 ng/mL) as a control. **(A)** The cells were collected and lysed. Ten micrograms of total protein was run on a reducing 4% bis-tris gel. α-SMA was detected by Western blotting using an α-SMA-specific primary antibody and the corresponding secondary antibody. Similarly, α-tubulin was also detected as a loading control. **(B)** Relative quantification of total α-SMA (normalized to α-tubulin) in NHBE cells. The data represent one independent experiment.

**Fig. S8.**
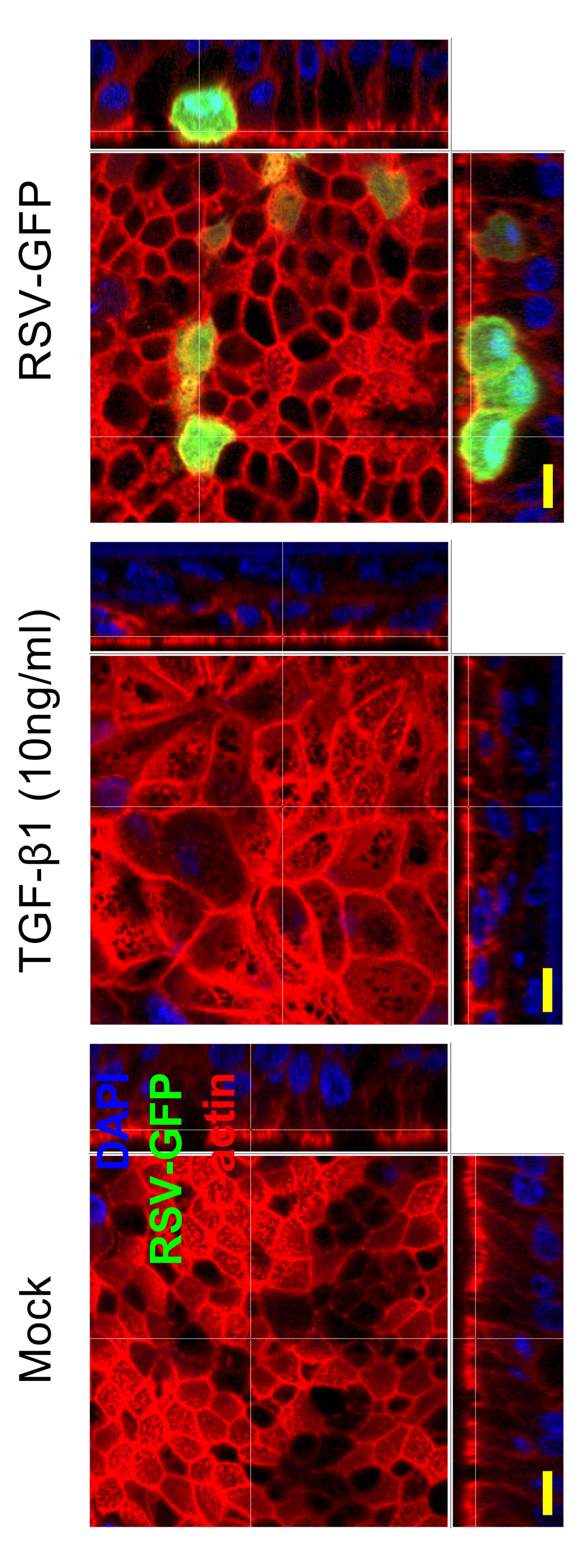
Differences in epithelial morphology between RSV infection and TGF-β1 treatment. Mock-infected or mock-treated (left), TGFβ1 (10 ng/ml)-treated (middle), or RSV-GFP-infected (MOI = 4, 6 dpi) (right) samples. RSV-infected cells were detected by the GFP signal. F-actin was visualized by rhodamine phalloidin (red). Images were captured with a 60× objective and then magnified (2.5×). The scale bar is 10 µm.

**Fig. S9.**
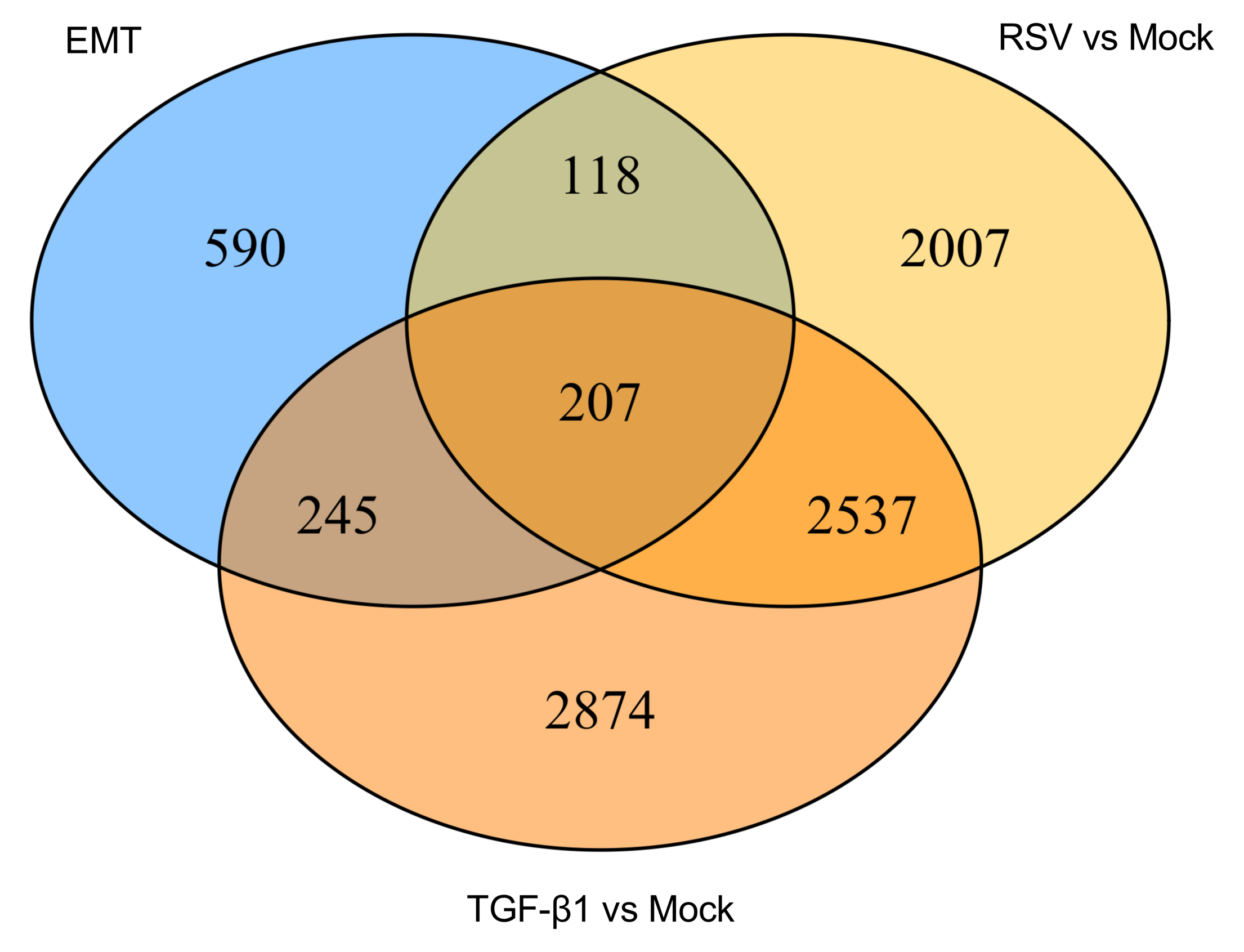
Common EMT genes regulated by either RSV infection, TGF-β1 treatment or both. A Venn diagram demonstrates that both RSV and TGF-β1 commonly modulate 207 EMT genes derived from the dbEMT database.

**Table S1.** Differentially expressed genes (DEGs).

**Table S2.** RSV-specific unique gene modulation in the ranking of DEGs

**Table S3.** TGF-β1-specific unique gene modulation in the ranking of DEGs **Table S4.** Enriched GO terms in RSV-specific DEGs

**Table S5.** Enriched GO terms shared by both TGF-β1-specific and RSV-specific cells.

**Table S6.** Enriched GO terms in TGF-β1-specific DEGs

**Table S7.** Genes shared by both TGF-β1-specific and RSV-specific effects

**Table S8.** Gene list of the EMT database dbEMT2.0.

**Table S9.** EMT-associated genes from dbEMT2.0 overlapping with both TGF-β1-specific and RSV-specific DEGs.

**Table S10.** Enriched GO terms linked to Clusters 1-3.

